# Gallbladder adenocarcinomas undergo subclonal diversification and selection from precancerous lesions to metastatic tumors

**DOI:** 10.1101/2022.03.31.486530

**Authors:** Minsu Kang, Hee Young Na, Soomin Ahn, Ji-Won Kim, Sejoon Lee, Soyeon Ahn, Ju Hyun Lee, Jeonghwan Youk, Haesook T. Kim, Kui-Jin Kim, Koung Jin Suh, Jun Suh Lee, Se Hyun Kim, Jin Won Kim, Yu Jung Kim, Keun-Wook Lee, Yoo-Seok Yoon, Jee Hyun Kim, Jin-Haeng Chung, Ho-Seong Han, Jong Seok Lee

**Author notes:** Co-corresponding authors: Ji-Won Kim, MD, PhD, Department of Internal Medicine, Seoul National University Bundang Hospital, Seoul National University College of Medicine, 82 Gumi-ro-173-beon-gil, Bundang-gu, Seongnam 13620, Korea; Soomin Ahn, MD, PhD, Department of Pathology and Translational Genomics, Samsung Medical Center, Sungkyunkwan University School of Medicine, 81 Irwon-ro, Gangnam-gu, Seoul 06351, Korea. These authors contributed equally to this work (Minsu Kang, Hee Young Na).

## Abstract

We aimed to elucidate the evolutionary trajectories of gallbladder adenocarcinoma (GBAC) using multi-regional and longitudinal tumor samples. Using whole-exome sequencing data, we constructed phylogenetic trees in each patient, and analyzed mutational signatures. A total of 11 patients including 2 rapid autopsy cases were enrolled. The most frequently altered gene in primary tumors was *ERBB2* (54.5%), followed by *TP53* (45.5%), and *FBXW7* (27.3%). Most mutations in frequently altered genes in primary tumors were detectable in concurrent precancerous lesions (biliary intraepithelial neoplasia, BilIN), but some of them were subclonal. Subclonal diversity was common in BilIN (n=4). However, among subclones in BilIN, a certain subclone commonly shrank in concurrent primary tumors. In addition, selected subclones underwent linear and branching evolution, maintaining subclonal diversity. In combined analysis with metastatic tumors (n=11), branching evolution was identified in 9 (81.8%) patients. Of these, 8 patients (88.9%) had a total of 11 subclones expanded at least 7-fold during metastasis. These subclones harbored putative metastasis-driving mutations in tumor suppressor genes such as *SMAD4*, *ROBO1*, and *DICER1*. In mutational signature analysis, 6 mutational signatures were identified: 1, 3, 7, 13, 22, and 24 (cosine similarity >0.9). Signatures 1 (age) and 13 (APOBEC) decreased during metastasis while signatures 22 (aristolochic acid) and 24 (aflatoxin) were relatively highlighted. Subclonal diversity arose early in precancerous lesions and the clonal selection was a common event during malignant transformation in GBAC. However, selected cancer clones continued to evolve and thus maintained subclonal diversity in metastatic tumors.

## Introduction

Gallbladder adenocarcinoma (GBAC) is a malignant neoplasm that has a high incidence rate in Chile, India, Poland, Pakistan, Japan, and Korea (1–3). Surgery is currently the only curative treatment modality for GBAC. However, because most patients are diagnosed at an advanced stage and thus inoperable, they receive palliative chemotherapy only. Therefore, the prognosis is poor with a median overall survival of only 11-15 months (3, 4).

The recent advancement of massively parallel sequencing technology has enabled us to deeply understand the genome of a variety of cancers. In GBAC, tumor suppressor genes such as *TP53*, *ARID1A,* and *SMAD4* and oncogenes such as *ERBB2* (*HER2*), *ERBB3*, and *PIK3CA* are significantly mutated (5–11). Of note, *ERBB2* amplification and overexpression occur in approximately 6.9-28.6% of GBAC (10, 12, 13) and may have therapeutic implications.

Cancer cells undergo clonal evolution by acquiring additional mutations and thus exhibit more aggressive phenotypes, including invasion and metastasis (14–16). Several large- scale studies have provided evidence of clonal evolution in some cancer types, including lung and kidney cancers (17, 18). However, there is no study so far that analyzed the patterns of clonal evolution from the initiation of carcinogenesis to distant metastasis in patients with GBAC.

This study aims to analyze the clonal evolutionary trajectories during carcinogenesis and metastasis of GBAC using multi-regional and longitudinal specimens including precancerous lesions (biliary intraepithelial neoplasia, BilIN), primary tumors, and metastatic tumors from patients who underwent biopsy, surgery, and rapid autopsy.

## Results

### Baseline characteristics of patients with GBAC

A total of 11 patients, including 2 rapid autopsy cases (GB-A1 and GB-A2) and 9 surgery cases (GB-S1 – 9), were enrolled in this study (**Table 1** and **Figure 1A**). There were 5 male and 6 female patients with the median age was 70 years (range, 59–75 years). Two patients had stage III and nine patients had IV disease at diagnosis. A total of 58 samples were analyzed, including 11 pairs of matched primary tumors and normal tissues, 6 concurrent BilIN, and 30 metastatic tumors: 15 were fresh frozen tissues and 43 were formalin-fixed paraffin-embedded (FFPE) tissues (**Supplementary Table 1**). The median number of filtered somatic SNVs and small indels was 61 (range, 12–241). The number of metastatic tumors in each patient ranged from 1 to 11.

**Figure 1.**
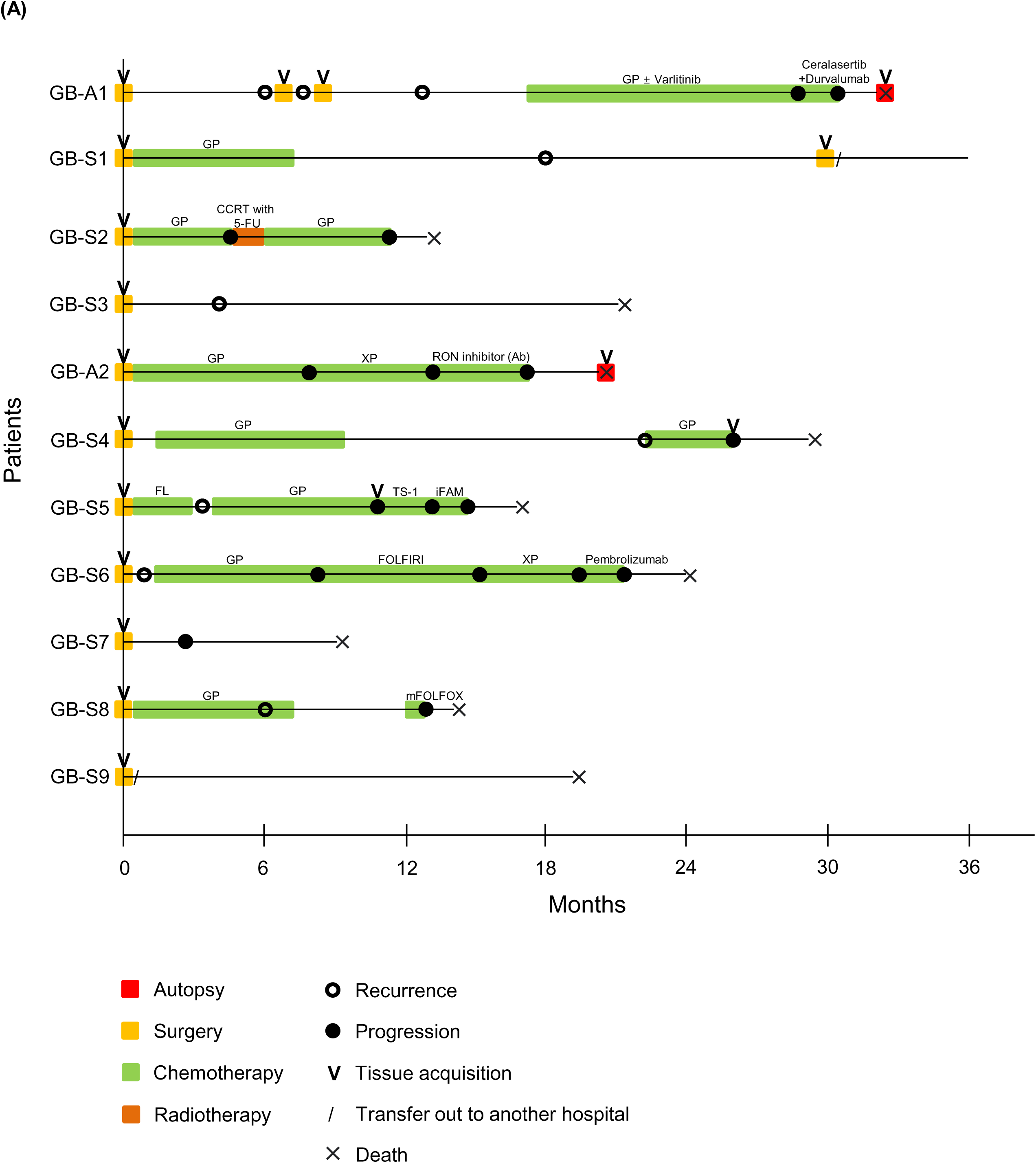
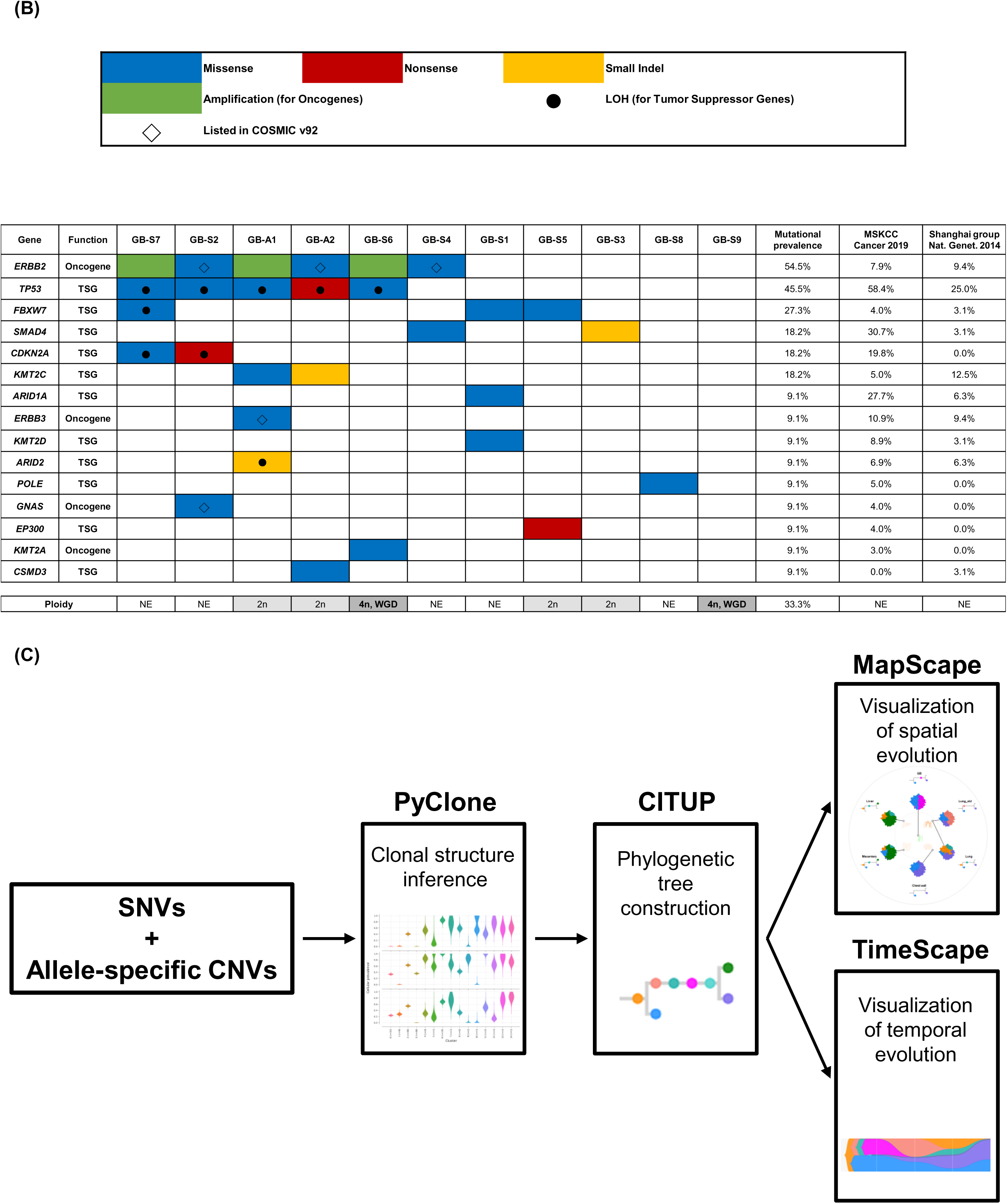
Clinical history of patients, mutational landscape, and study workflow. **(A**) The swimmer plot showing the clinical history of 11 patients. **(B**) The mutational landscape of 11 primary tumors visualized according to the prevalence and compared with the results of two previous studies on GBAC. **(C)** Workflow for constructing clonal evolution trajectories using multiple tumor samples. Adj, adjuvant; CCRT, concurrent chemoradiotherapy; CNVs, copy number variations; FL, 5- fluorouracil + leucovorin; FOLFIRI, 5-fluorouracil + leucovorin + irinotecan; GB, gallbladder; GP, gemcitabine + cisplatin; iFAM, infusional 5-fluorouracil + doxorubicin + mitomycin-C; LOH, loss-of-heterozygosity; mFOLFOX, modified 5-fluorouracil + leucovorin + oxaliplatin; NE, not evaluable; SNVs, single nucleotide variants; TSG, tumor suppressor gene; WGD, whole genome doubling; XP, capecitabine + cisplatin; 5-FU, 5-fluorouracil.

**Table 1.** Baseline characteristics of 11 patients with GBAC.

| **Patient ID\*** | **Sex** | **Age at diagnosis** | **ECOG PS**<br>**at diagnosis** | **Stage**<br>**at diagnosis** | **Differentiation** |
| --- | --- | --- | --- | --- | --- |
| GB-A1 | F | 70 | 1 | IV | PD |
| GB-S1 | F | 66 | 1 | IV | MD |
| GB-S2 | F | 66 | 1 | IV | MD |
| GB-S3 | F | 75 | 1 | III | MD |
| GB-A2 | M | 61 | 1 | IV | MD |
| GB-S4 | M | 72 | 1 | IV | MD |
| GB-S5 | F | 70 | 1 | IV | PD |
| GB-S6 | M | 70 | 0 | IV | MD |
| GB-S7 | F | 74 | 1 | IV | PD |
| GB-S8 | M | 67 | 1 | III | WD |
| GB-S9 | M | 59 | 0 | IV | MD |

\* In patient ID, “A” indicates autopsy cases whereas “S” indicates surgery cases.
GBAC, gallbladder adenocarcinoma; ECOG PS, Eastern Cooperative Oncology Group performance status; M, male; F, female; PD, poorly differentiated; MD, moderately differentiated; WD, well-differentiated.

### Mutational landscape and ploidy of primary tumors

The mutational landscape of the 11 primary tumors was analyzed and compared with previous literature (**Figure 1B**) (10, 11). The most frequently altered gene was *ERBB2* (54.5%), followed by *TP53* (45.5%) and *FBXW7* (27.3%). *ERBB2* amplification was defined as a copy number ≥ 6 (19). Of the six *ERBB2* alterations, three were amplification and the other three were missense mutations listed in the COSMIC (Catalogue of Somatic Mutations in Cancer) v92 database. All *TP53* mutations found in 5 patients were accompanied by loss-of-heterozygosity (LOH). Three *ERBB2* SNVs and one *ERBB3* SNV were pathogenic or likely pathogenic in the ClinVar database (20).

Ploidy was analyzed in 15 tumors of 6 patients with purity > 0.4 because ploidy estimation was inaccurate when tumor purity is ≤ 0.4 (21). In 2 patients (GB-S6 and GB-S9, 33.3%), WGD was detected in both the primary and metastatic tumors (**Figure 1B**). In GB-S5 patient, WGD was found in distant metastasis, but not in primary GBAC.

### Somatic mutations developed at the precancerous stage

Multi-regional distribution and longitudinal evolution of clones were analyzed using PyClone (**Figure 1—figure supplement 1** and **Supplementary File 1**) (22) and CITUP (23) and then visualized using MapScape and TimeScape (24), respectively (**Figure 1C**). Among 6 patients having concurrent BilIN tissues, two patients were excluded from the further analysis because of low tumor purity in one patient and different mutational profiles between BilIN and primary GBAC in the other patient, suggesting different origins of the two tumors (**Figure 1—figure supplement 2**).

Most mutations in frequently altered genes in GBAC existed at the BilIN stage (10 of 13, 76.9%), but some of them were subclonal. Mutations in representative oncogenes and tumor suppressor genes were described (**Figure 2**). In GB-A1 (**Figure 2A** and **B**), *TP53* C100Y with LOH, *KMT2C* R909K*, ERBB3* G284R*, ARID2* F575fs with LOH, and *CTNNB1* S37F with LOH were observed clonally in BilIN. In GB-S1 (**Figure 2C**), *SAMD9* E612K was clonal whereas *FBXW7* S18C*, ARID1A* S382N, and *NF1* R440X were subclonal. In GB-S2 (**Figure 2D**), considering the cellular prevalence of mutations, it is speculated that the mutations developed in the order of *CDKN2A* R58fs with LOH, *TP53* H82R with LOH, *ERBB2* V777L, and *GNAS* G306R during carcinogenesis. In GB-S3 (**Figure 2E**), *ERBB3* E332K, which was listed in COSMIC v92, was dominant in BilIN, while *SMAD4* 539_540del was not detected.

**Figure 2.**
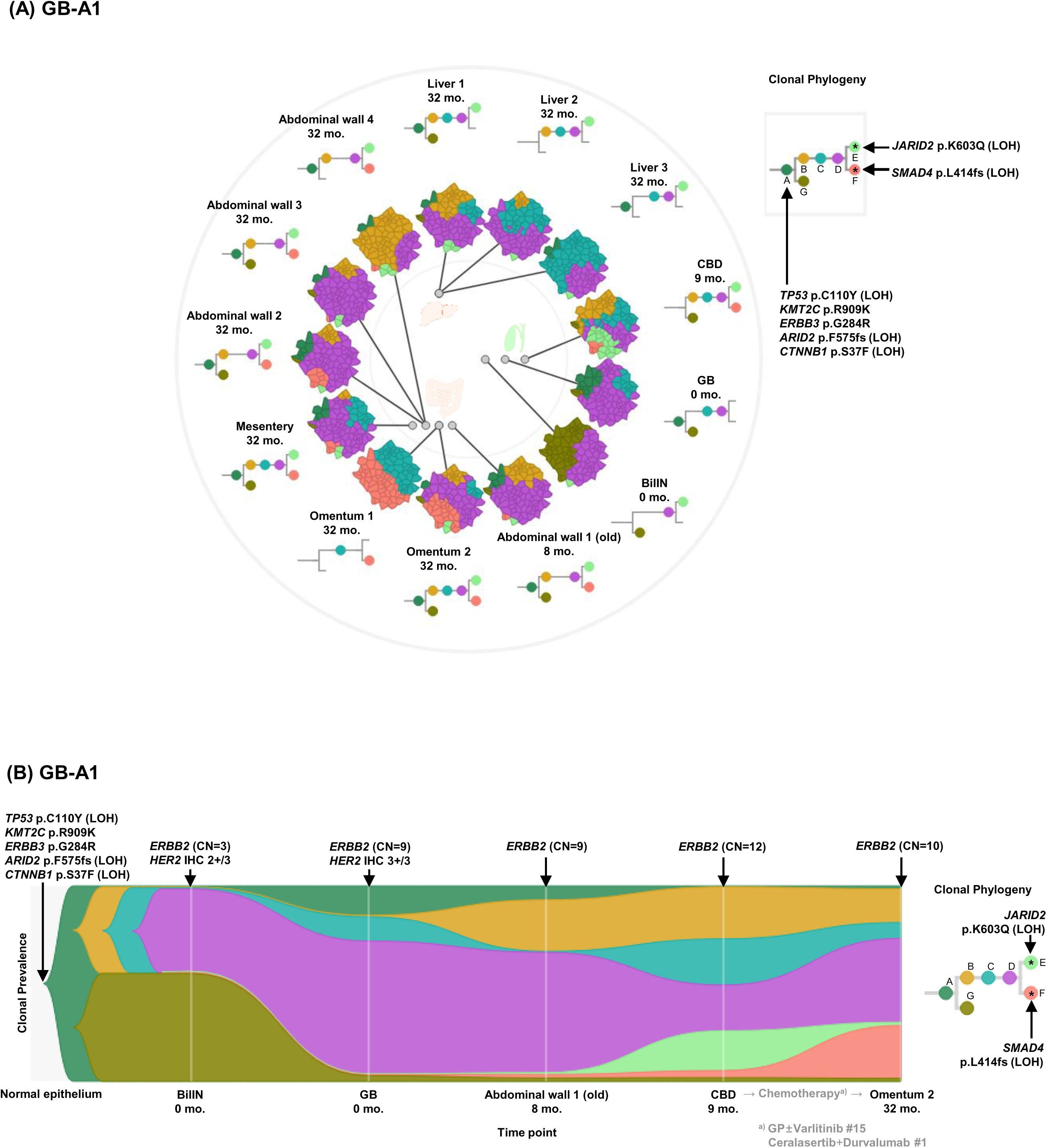
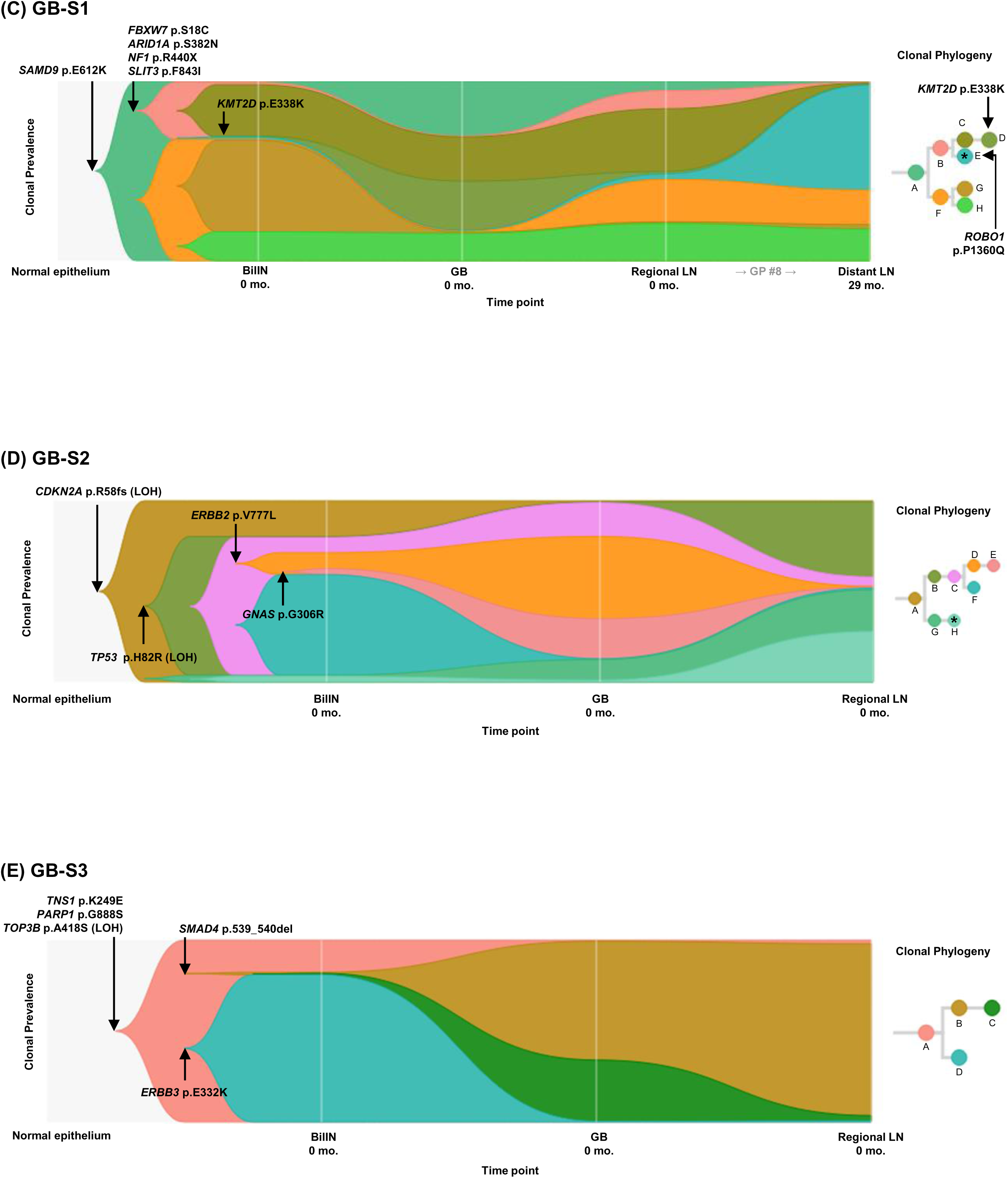
Spatial and temporal clonal evolution of 4 patients with GBAC whose precancerous BilIN tissues were analyzed. **(A-E)** The most probable phylogenetic trees were constructed for GB-A1 **(A,** MapScape), GB- A1 **(B,** TimeScape), GB-S1 **(C,** TimeScape), GB-S2 **(D,** TimeScape), and GB-S3 **(E,** TimeScape) using PyClone and CITUP. In MapScape visualization, spatially distinct tumor samples were indicated with an anatomical image. Colors represent distinct clones and clonal prevalences per site were proportional to the corresponding-colored area of the cellular aggregate representation. In TimeScape visualization, clonal prevalences (vertical axis) were plotted across timepoints (horizontal axis) for each clone (colors). Asterisks (*) in the clonal phylogenetic tree denote subclones that constituted <5% in the primary tumor and expanded more than 7-fold in the metastatic tumor. Notable events were marked with arrows. The time from diagnosis of GBAC to tissue acquisition was indicated under the sample name. Chemotherapy history was indicated in gray color, where ‘#’ represents the number of chemotherapy cycles. BilIN, biliary intraepithelial neoplasia; CBD, common bile duct; CN, copy number; GB, gallbladder; GP, gemcitabine + cisplatin; IHC, immunohistochemistry; LN, lymph node; LOH, loss-of-heterozygosity.

### Subclonal diversity and ‘selective sweep’ phenomenon during the early stage of *carcinogenesis*

Branching evolution and subclonal diversity were commonly observed in BilIN of the 4 patients (**Figure 2A**-**E**). When compared with the concurrent primary tumors, one subclone commonly shrank in the primary tumors, while the other subclones that acquired additional mutations relatively expanded in the primary tumors, suggesting a selective sweep phenomenon (25). Selected subclones underwent linear and branching evolution, and thus subclonal diversity was maintained after the BilIN stage. In GB-A1 (**Figure 2B**), clone A underwent branching evolution into B and G, and clone B linearly evolved into C then D. Clone D, which acquired additional mutations, increased from 42.7% to 68.1% while clone G decreased from 55.7% to 2.8%. In GB-S1 (**Figure 2C**), clone D that acquired *KMT2D* E338K increased from 0.1% to 28.1%, while clone G decreased from 51.4% to 0%. In GB-S2 (**Figure 2D**), clone D that acquired *ERBB2* V777L, clone E that acquired *GNAS* G306R, and clone G increased from 9.0%, 3.5%, and 0% to 45.5%, 22.3%, and 11.4%, respectively. In contrast, clone F decreased from 55.8% to 0.6%. In GB-S3 (**Figure 2E**), clone B that acquired *SMAD4* 539_540del increased from 0.1% to 65.6%, while clone D containing *ERBB3* E332K decreased from 81.1% to 0.6%.

### Evolutionary trajectories and expansion of subclones during regional and distant metastasis

Combined analysis of regional and distant metastatic tumors revealed branching evolution in 9 patients (81.8%) and linear evolution in 2 patients (18.2%). Of the 9 patients with branching evolution, eight (88.9%) had a total of 11 subclones expanded at least 7-fold in the regional or distant metastasis stage (**Table 2**). Of these 11 subclones, putative driver mutations in tumor suppressor genes were described in 8 subclones. In GB-A1 (**Figure 2A** and **B**), clone E, which acquired *JARID2* (26) K603Q with LOH, increased from 0.2% to 20.5% during common bile duct (CBD) metastasis. In addition, clone F, which acquired *SMAD4* (10, 17, 27, 28) L414fs with LOH, expanded from 0.4% to 49.8% during omentum 1 metastasis and to 27.0% during omentum 2 metastasis. In GB-S1 (**Figure 2C**), clone B harboring *SLIT3* F843I mutation evolved into E by additionally acquiring *ROBO1* P1360Q mutation (29). In GB-A2 (**Figure 3A**-**C**), clone F, which acquired *PRKCD* (30, 31) I153L, expanded from 0.3% to 73.1% and 68.3% during metastasis to liver and mesentery, respectively. In addition, clone G acquired *DICER1* (32) T519A and expanded from 2.8% to 39.9% and 61.5% during metastasis to lung and chest wall, respectively. In the remaining 6 subclones in 5 patients (**Figure 2D** and **3D-I**), mutations in *FBXW2* (33), *KIAA0100* (34), *CSMD2* (34), and *OSCP1* (35) genes were observed.

**Figure 3.**
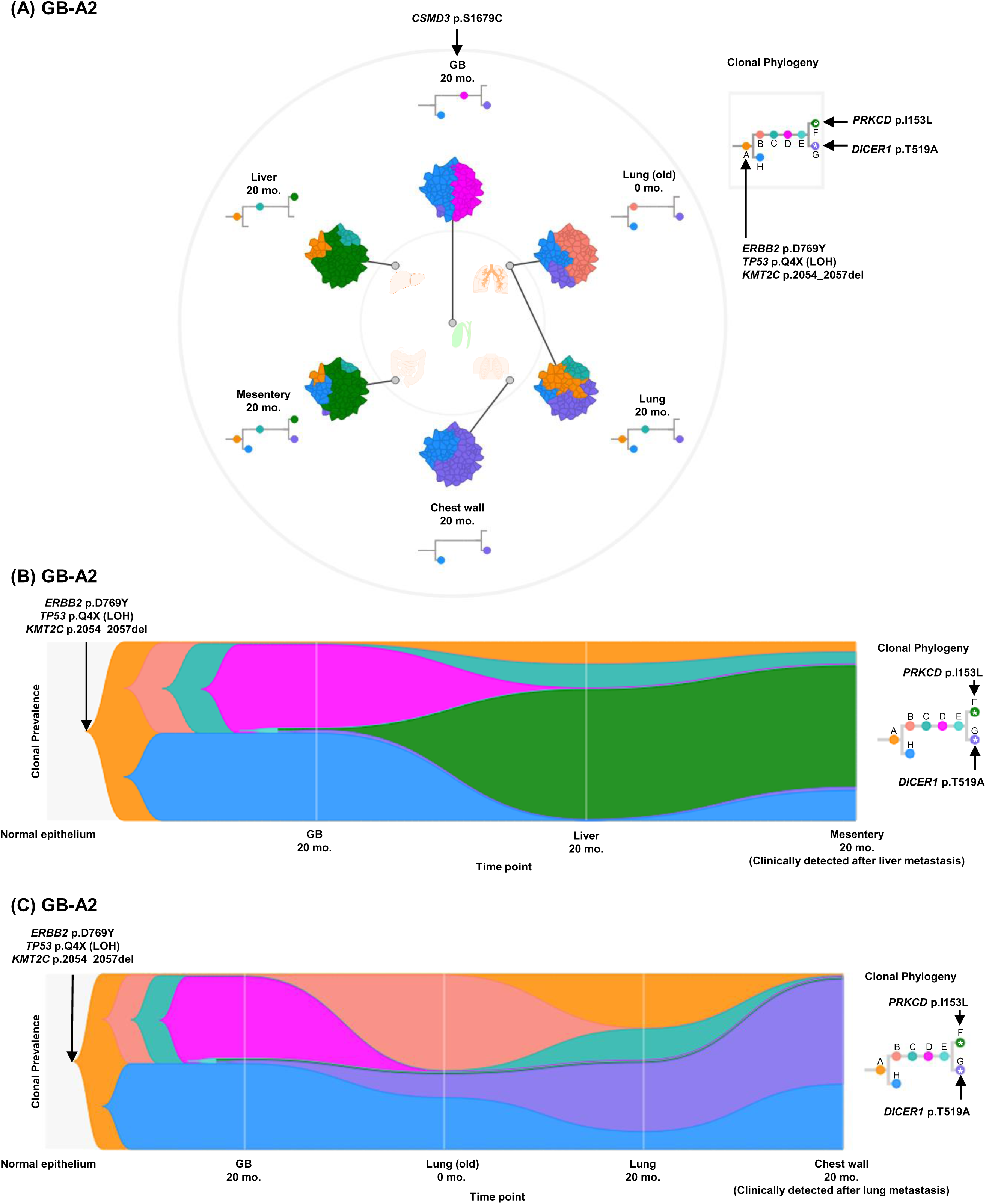
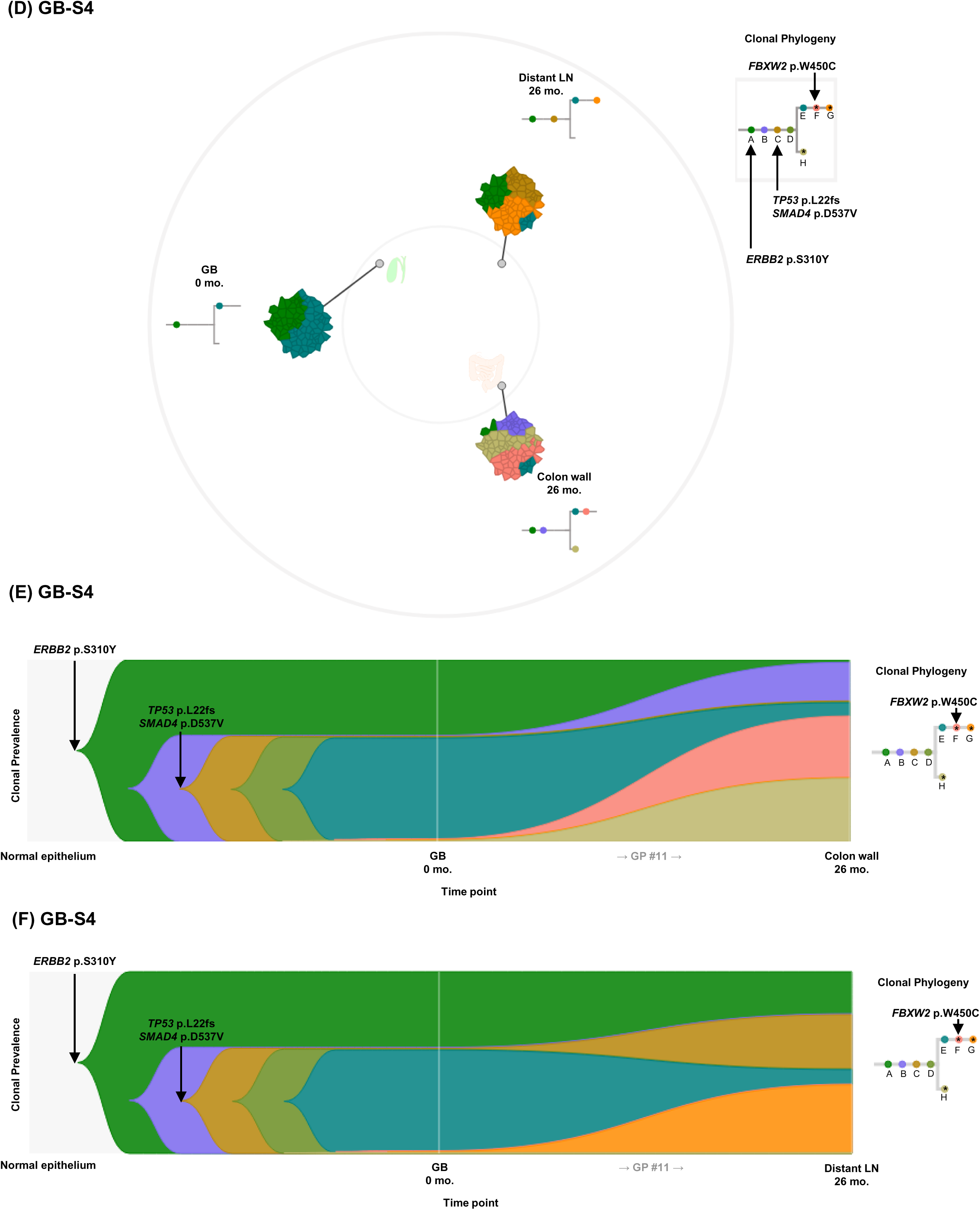
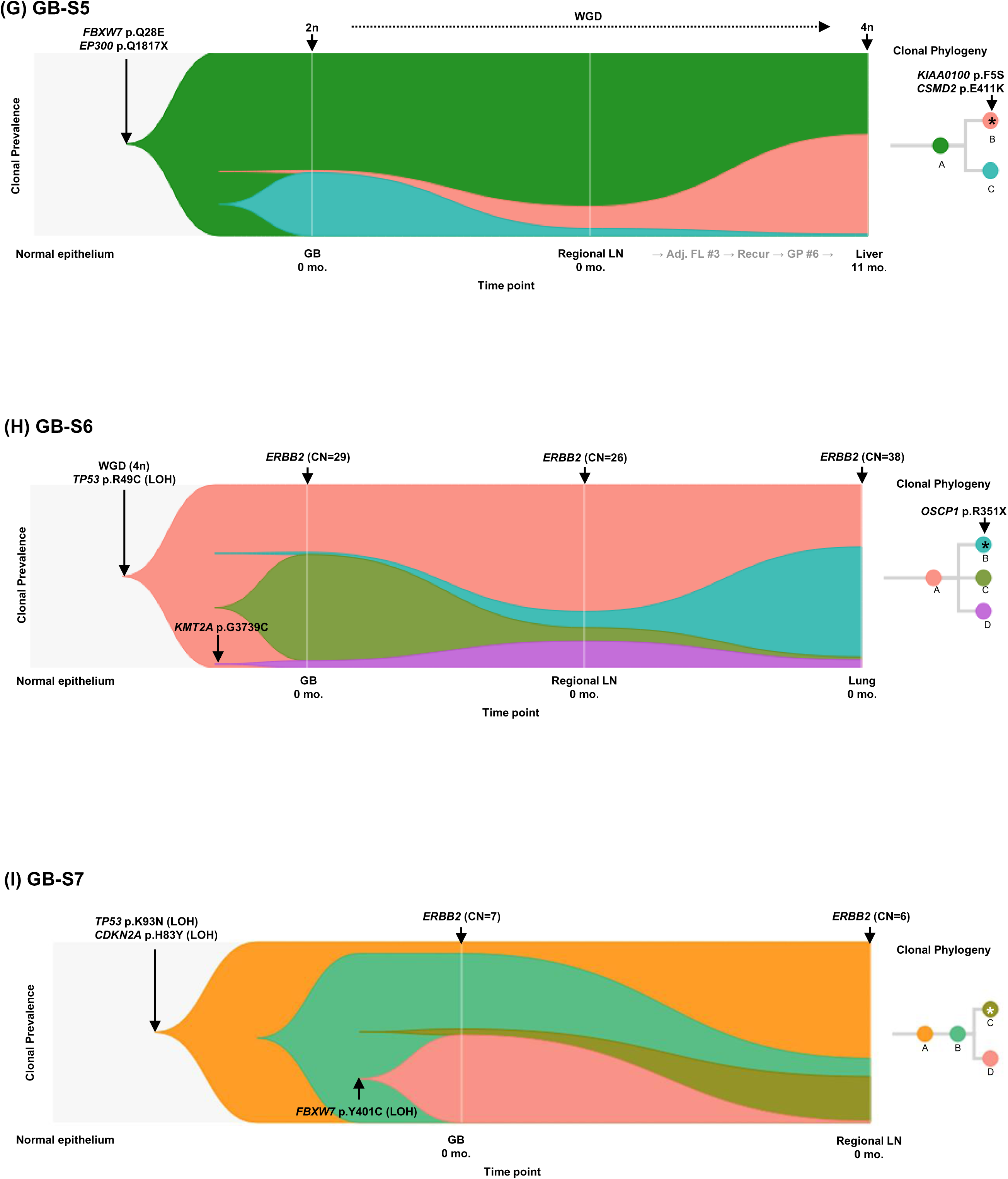
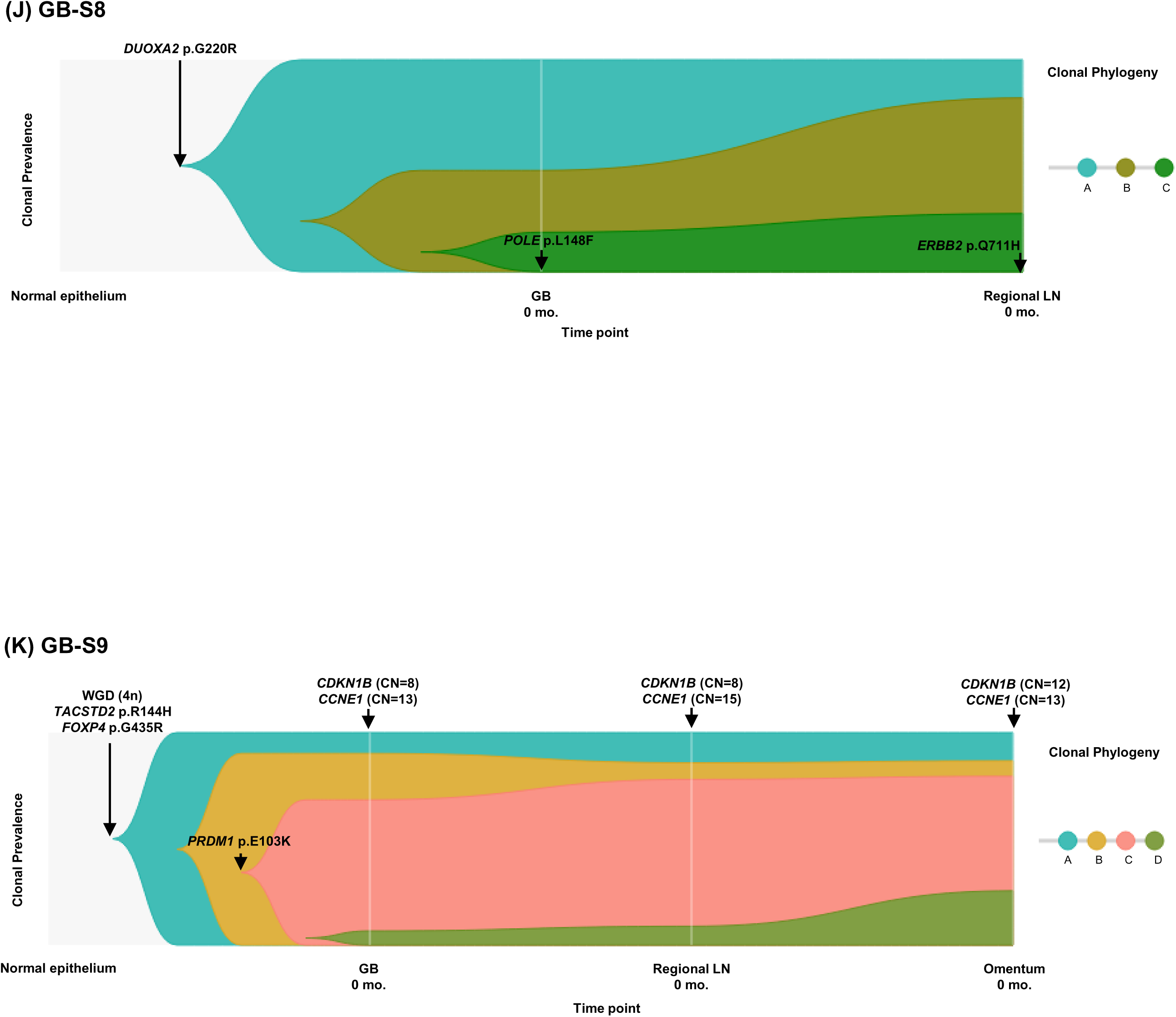
Spatial and temporal clonal evolution of additional 7 patients with GBAC. **(A-K)** The most probable phylogenetic trees were constructed for GB-A2 (**A,** MapScape), GB- A2 (**B,** TimeScape), GB-A2 (**C,** TimeScape), GB-S4 (**D,** MapScape), GB-S4 (**E,** TimeScape), GB-S4 (**F,** TimeScape), GB-S5 (**G,** TimeScape), GB-S6 (**H,** TimeScape), GB-S7 (**I,** TimeScape), GB-S8 (**J,** TimeScape), and GB-S9 (**K,** TimeScape) using PyClone and CITUP. In MapScape visualization, spatially distinct tumor samples were indicated with an anatomical image. Colors represent distinct clones and clonal prevalences per site were proportional to the corresponding-colored area of the cellular aggregate representation. In TimeScape visualization, clonal prevalences (vertical axis) were plotted across timepoints (horizontal axis) for each clone (colors). Asterisks (*) in the clonal phylogenetic tree denote subclones that constituted <5% in the primary tumor and expanded more than 7-fold in the metastatic tumor. Notable events were marked with arrows. The time from diagnosis of GBAC to tissue acquisition was indicated under the sample name. Chemotherapy history was indicated in gray color, where ‘#’ represents the number of chemotherapy cycles. Adj, adjuvant; BilIN, biliary intraepithelial neoplasia; CN, copy number; FL, 5-fluorouracil + leucovorin; GB, gallbladder; GP, gemcitabine + cisplatin; IHC, immunohistochemistry; LN, lymph node; LOH, loss-of-heterozygosity; WGD, whole genome doubling.

**Table 2.** List of subclones expanding during metastasis.

| Patient ID | Subclone | No. of mutations | Putative driver mutations | Clonal prevalence in primary tumor | Clonal prevalence in metastasis | Metastatic organ |
| --- | --- | --- | --- | --- | --- | --- |
| GB-A1 | E | 121 | *JARID2* p.K603Q (LOH) | 0.2% | 20.5% | CBD |
|  | F | 54 | *SMAD4* p.L414fs (LOH) | 0.4% | 49.8% | Omentum 1 |
|  |  |  |  |  | 27.0% | Omentum 2 |
| GB-S1 | E | 6 | *ROBO1* p.P1360Q | 0.0% | 59.2% | Distant LN |
| GB-S2 | H | 6 | - | 1.2% | 28.1% | Regional LN |
| GB-A2 | F | 14 | *PRKCD* p.I153L | 0.3% | 73.1% | Liver |
|  |  |  |  |  | 68.3% | Mesentery |
|  | G | 3 | *DICER1* p.T519A | 2.8% | 39.9% | Lung |
|  |  |  |  |  | 61.5% | Chest wall |
| GB-S4 | F | 32 | *FBXW2* p.W450C | 0.0% | 34.1% | Colon wall |
|  | H | 36 | - | 0.0% | 34.9% | Distant LN |
| GB-S5 | B | 28 | *KIAA0100* p.F5S<br>*CSMD2* p.E411K | 1.2% | 54.4% | Liver |

| **Patient ID** | **Subclone** | **No. of mutations** | **Putative driver mutations** | **Clonal prevalence in primary tumor** | **Clonal prevalence in metastasis** | **Metastatic organ** |
| --- | --- | --- | --- | --- | --- | --- |
| GB-S6 | B | 7 | *OSCP1* p.R351X | 1.1% | 60.2% | Lung |
| GB-S7 | C | 5 | - | 3.3% | 24.7% | Regional LN |

Putative driver mutations in tumor suppressor genes are indicated, and a full list of mutated genes is specified in Supplementary File 1.
GB, gallbladder; LN, lymph node; CBD, common bile duct.

In GB-S8 and GB-S9, linear evolution was identified (**Figure 3J** and **K**). In GB-S9 (Figure 3K), which did not harbor any mutation in frequently altered genes in GBAC (**Figure 1B**), amplification of cell cycle-related oncogenes *CDKN1B* and *CCNE1* was uniformly observed from primary GBAC to regional and distant metastatic tumors.

### Polyclonal metastasis and intermetastatic heterogeneity

The metastatic lesions were uniformly polyclonal. In GB-A1, GB-A2, and GB-S4, which contained two or more distant metastatic lesions, the clonal compositions of metastatic lesions were heterogeneous. However, metastatic lesions in one organ or adjacent organs showed similar clonal compositions. In GB-A1 (**Figure 2A**), abdominal wall 1-4 did not contain clone C, and liver 1-3 did not contain clone F. In addition, omentum 1-2 had a high prevalence of clone F of over 27.0%. Notably, clone F, which was not found in both BilIN and primary GBAC, first appeared in abdominal wall 1 (old) 8 months later, and then was observed in CBD 9 months later and omentum 1-2, mesentery, and abdominal wall 2-4 lesions 32 months later. In GB-A2 (**Figure 3A**-**C**), the proportion of clone F was specifically high in liver and mesentery lesions, while the proportion of clone G was specifically high in lung and chest wall lesions. In clinical information of the GB-A2 patient, mesentery and chest wall metastases developed later than liver and lung metastases. In GB-S4 (**Figure 3D**-**F**), proportions of clone G and H were specifically high in distant LN and colon wall metastasis, respectively.

### Mutational signatures during clonal evolution

We compared our mutational signature analysis results (**Figure 4—figure supplement 1**) with those of MSKCC and Shanghai datasets (10, 11), according to COSMIC Mutational Signatures v2 (36). While MSKCC and Shanghai datasets consist merely of primary tumors, our dataset includes precancerous and metastatic lesions. Signatures 1 (age), 3 (DNA double-strand break-repair), and 13 (APOBEC) were commonly dominant in all three datasets, while signatures 22 (aristolochic acid) and 24 (aflatoxin) were exclusive in our dataset and signatures 2 (APOBEC), 9 (DNA polymerase η), and 29 (tobacco chewing) were found only in MSKCC or Shanghai datasets (cosine similarity > 0.9 in all datasets) (**Figure 4A**). We then analyzed the mutations according to the developmental stages of cancer (**Figure 4B**) as follows: (1) BilIN, (2) primary GBAC, (3) regional LN metastasis, and (4) distant metastasis. In distant metastasis tumors, signatures 1, 7, and 13 relatively decreased, while signatures 22 and 24 increased, compared with the other 3 tumors. Next, we analyzed the mutations according to the timing of development during clonal evolution (**Figure 4C**): (1) early carcinogenesis (*i.e.* clone A in **Figure 2** and **3**), (2) late carcinogenesis (*i.e.* subclones which were not categorized in early carcinogenesis or metastasis), and (3) metastasis (*i.e.* subclones expanding during metastasis [Table 2]). At metastasis phase, signatures 1 and 13 decreased while signatures 22 and 24 increased compared with early and late carcinogenesis.

**Figure 4.**
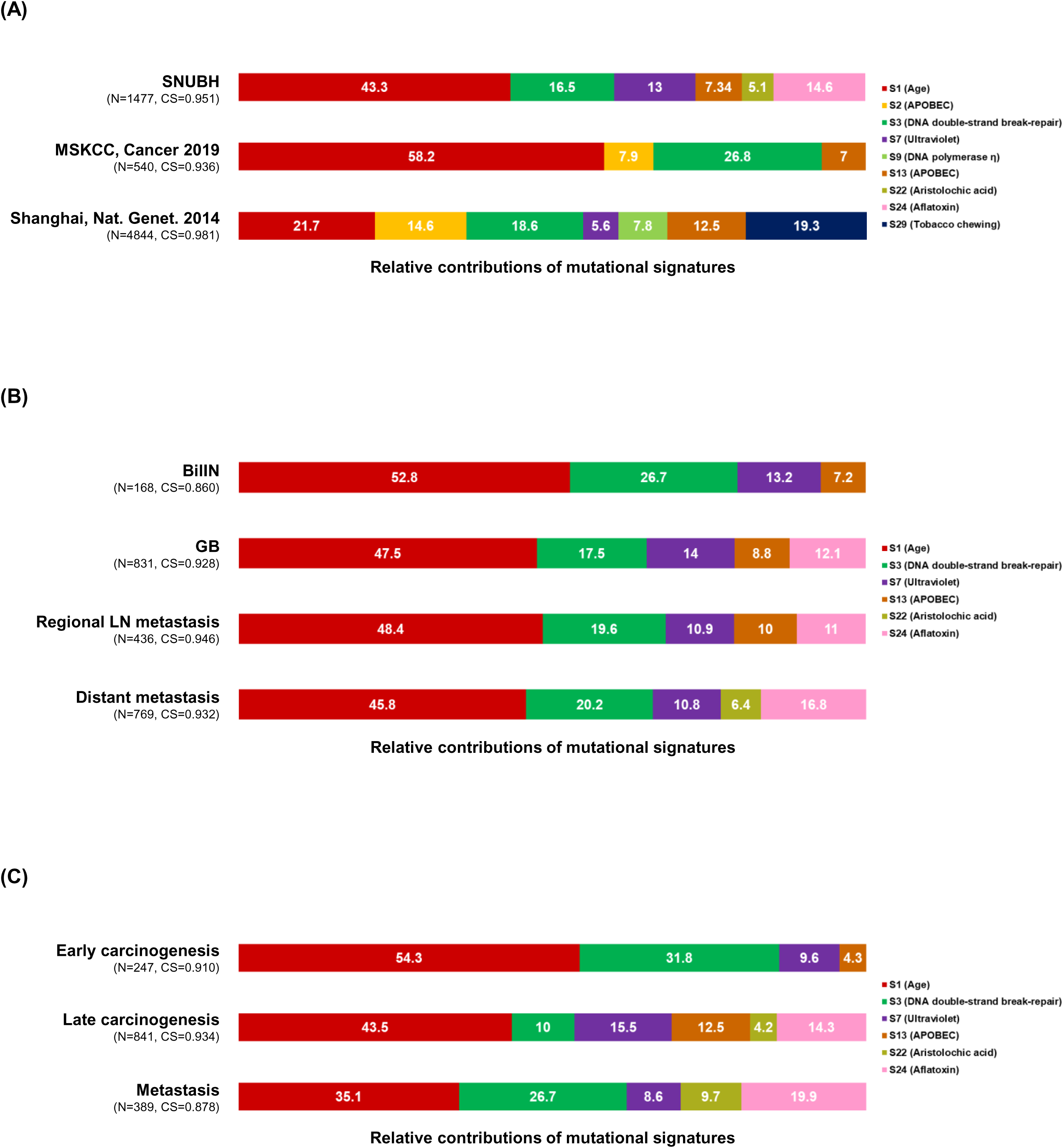
Mutational signature analysis. **(A-C**) The 100% stacked bar plots comparing the proportions of known COSMIC Mutational Signatures v2 within our dataset and two public (MSKCC and Shanhai) datasets (**A**), each category split according to the developmental stages of cancer (**B**) and each category split according to the timing of development during clonal evolution (**C**). The total number of mutations (N) and cosine similarity (CS) values of each category were noted. BilIN, biliary intraepithelial neoplasia; COSMIC, catalogue of somatic mutations in cancer; GB, gallbladder; LN, lymph node.

### *ERBB2* amplification during clonal evolution

In GB-A1 (**Figure 5**), *ERBB2* copy number (**Figure 5A** and **B**) was 3 and 9 in concurrent BillN and primary tumors, respectively. The increased copy number of *ERBB2* was maintained after distant metastasis (**Figure 2B**). *HER2* (=*ERBB2*) SISH (**Figure 5C** and **D**) was conducted to evaluate the subclonal distribution of *ERBB2* amplification at the single BillN cell level. *ERBB2* copy number per cell ranged from 1 to 5 in BilIN, and from 2 to 14 in primary GBAC. *ERBB2*/CEP17 ratio was 2.48 and 6.00 in BilIN and primary GBAC, respectively (**Figure 5E**). In addition, *HER2* IHC (**Figure 5F** and **G**) was carried out to evaluate whether the *ERBB2* amplification was correlated to *HER2* protein expression levels on the membrane of tumor cells. The *HER2* IHC results were 2+/3 in the BillN and 3+/3 in primary GBAC. Therefore, *ERBB2* amplification might have initiated from the precancerous stage and further progressed during the malignant transformation of BilIN, resulting in increased *HER2* protein expression levels. In GB-S6 and GB-S7 (**Figure 3H** and **I**), *ERBB2* amplification was maintained in primary tumors and regional and distant metastatic tumors.

**Figure 5.**
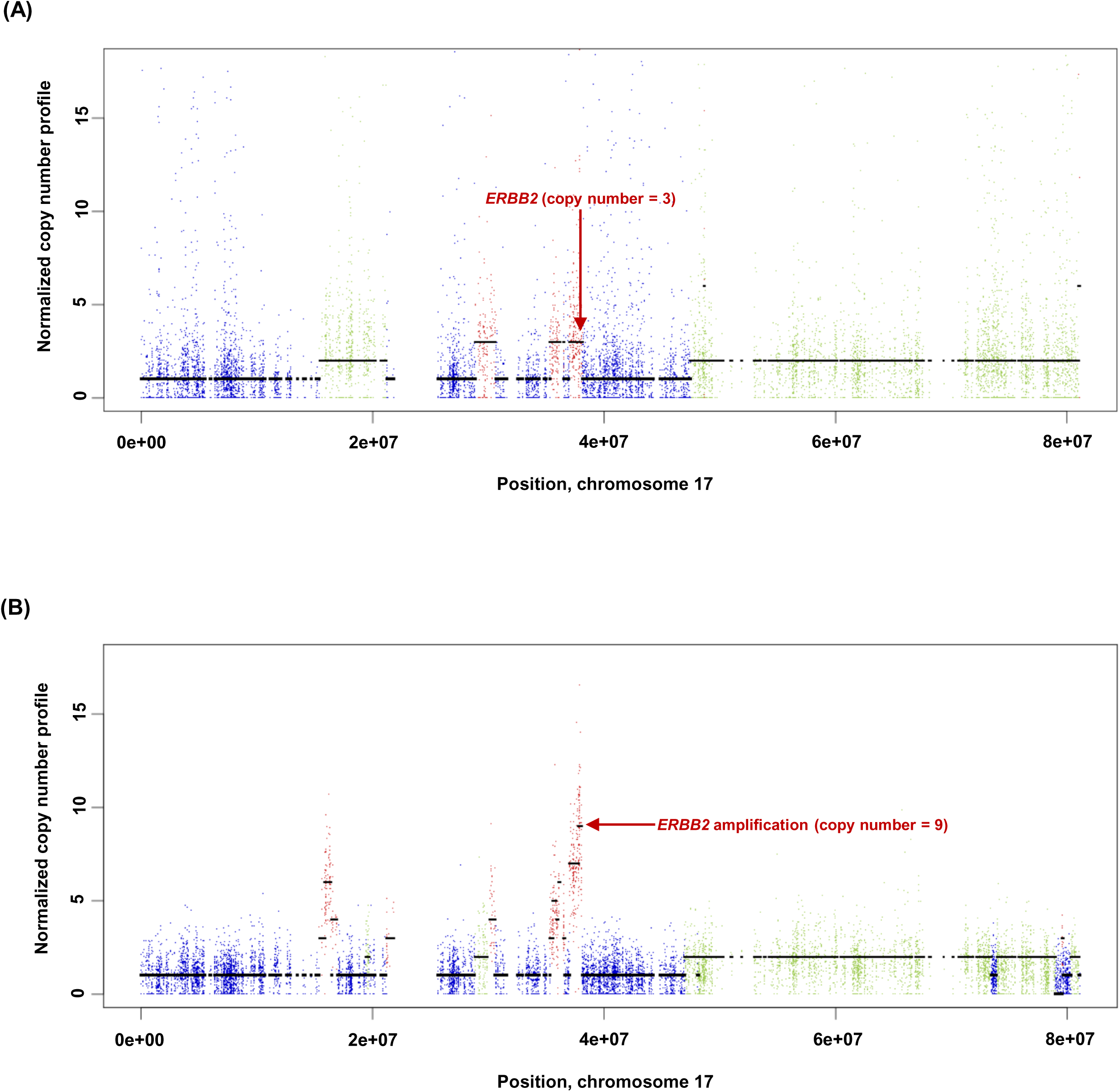
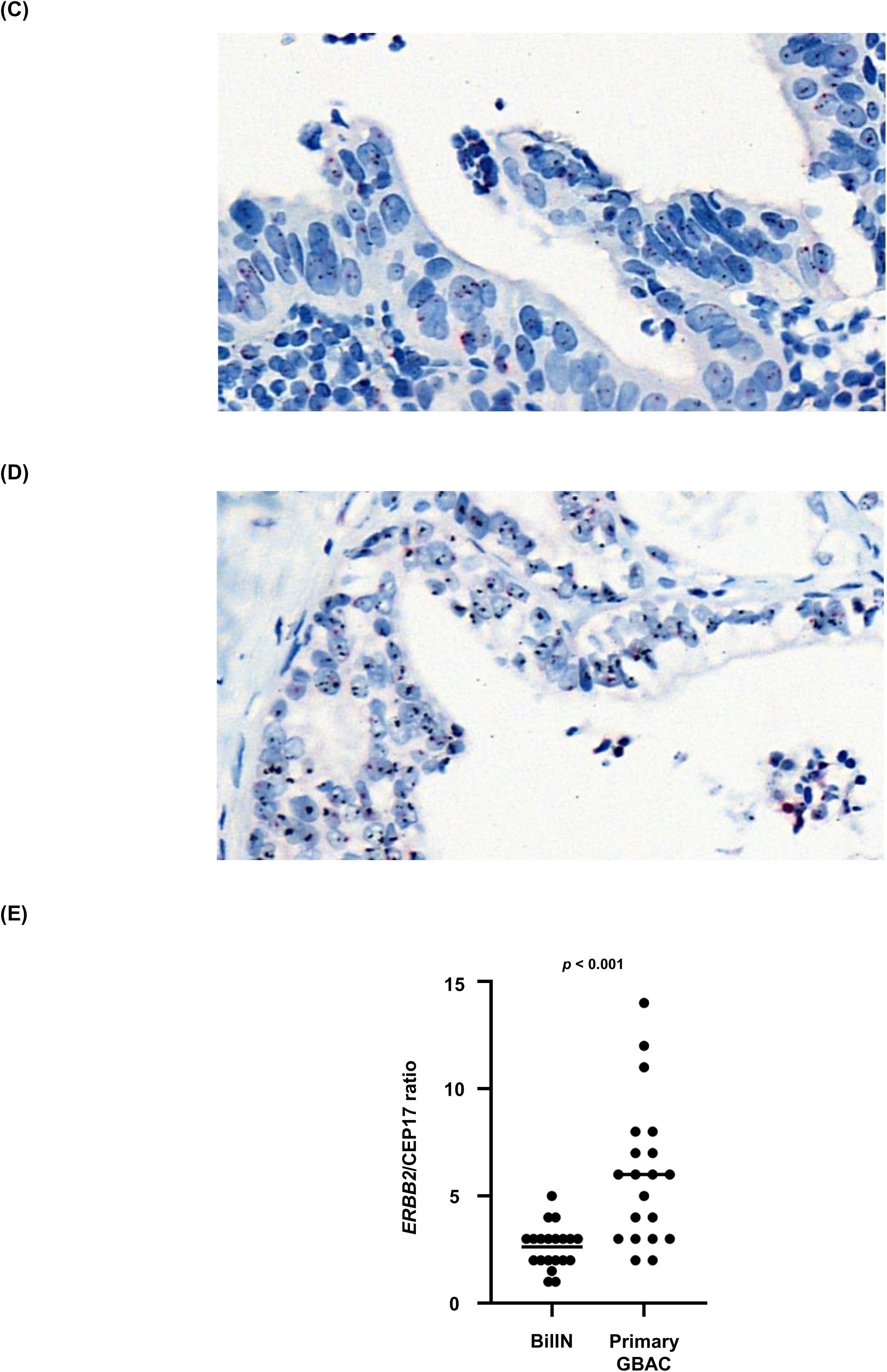
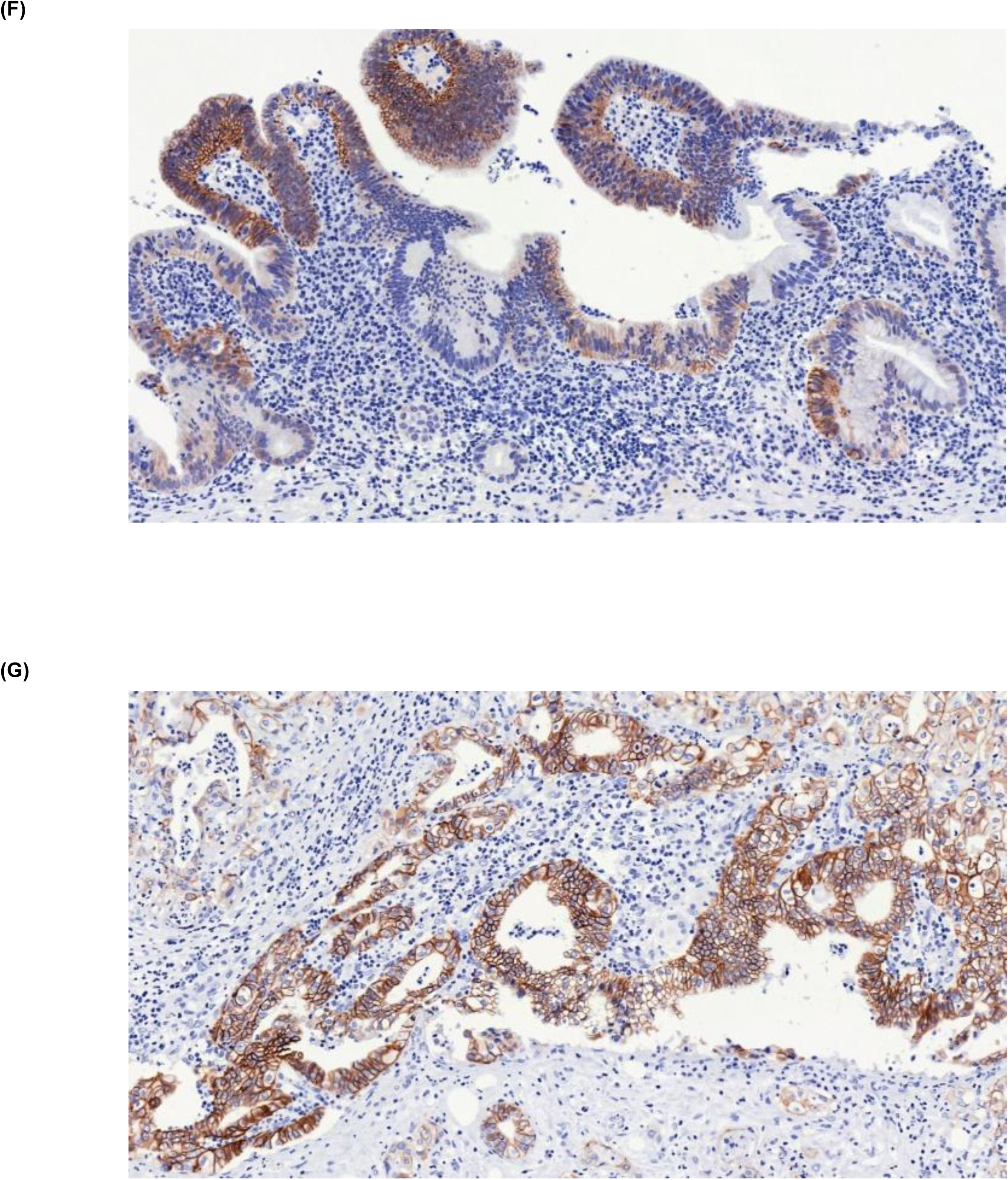
*ERBB2* copy number variation during neoplastic transformation of BilIN in GB-A1. **(A-G)** *ERBB2* gene amplification (**A** and **B**), *HER2* SISH (**C** and **D**), and *HER2* IHC (**F** and **G**) were compared between BilIN (**A**, **C,** and **F**) and primary GBAC (**B**, **D**, and **G**) samples and the mean *ERBB2*/CEP17 ratio of BilIN and GB-A1 samples were compared by using the Wilcoxon rank-sum test (**E**). The copy number was 3, IHC result was 2+/3, and *ERBB2*/CEP17 ratio was 2.48 in BilIN whereas the copy number was 9, IHC result was 3+/3, and the mean *ERBB2*/CEP17 ratio was 6.00 in primary GBAC. BilIN, biliary intraepithelial neoplasia; GBAC, gallbladder adenocarcinoma.

## Discussion

To the best of our knowledge, this is the first study to investigate clonal evolution from precancerous lesions to metastatic tumors in patients with GBAC. In this study, evolutionary trajectories of GBAC were inferred using multi-regional and longitudinal WES data from precancerous lesions (BilIN) to primary and metastatic tumors. Based on these results, we derived comprehensive models of carcinogenesis and metastasis in GBAC.

In our analysis of carcinogenesis, we discovered three common themes. First, most mutations in frequently altered genes in primary GBAC are detected in concurrent BilIN (10 of 13, 76.9%), but some of them are subclonal. Second, branching evolution and subclonal diversity are commonly observed at the BilIN stage. Third, one subclone in BilIN commonly shrinks in the primary tumors, while the other subclones undergo linear and branching evolution, maintaining subclonal diversity after the BilIN stage. A previous study in colorectal cancer by Vogelstein and colleagues demonstrated a stepwise carcinogenesis model from the precancerous lesion, adenoma, to invasive carcinoma by the accumulation of mutations, called the adenoma-carcinoma sequence (37). In addition, recent studies on esophageal squamous cell carcinoma have reported that not only dysplasia but also histologically normal epithelia frequently harbor cancer-driving mutations (38, 39). A recent study on GBAC reported that *CTNNB1* mutation was frequently observed (5 out of 11) when BilIN and primary GBAC coexist (9). In our BilIN analysis, *CTNNB1* S37F with LOH was observed in 1 (GB-A1) of 4 patients.

In the analysis of metastasis, the following three phenomena are observed. First, subclonal expansion is frequent (8 of 11 patients, 72.7%) and some subclones expand substantially in metastatic tumors, leading to increased subclonal diversity. Previous studies suggest that subclonal diversity increases through branching evolution during progression and metastasis (40–43). Second, metastases are polyclonal but metastatic lesions in one organ or adjacent organs show similar clonal compositions. Previously, it was thought that metastasis is initiated by the migration of a single cell to another organ (44). However, recent data suggest that polyclonal seeding occurs due to the migration of a cluster of cancer cells (14, 45–47). Third, we found evidence of metastasis-to-metastasis spread. In recent studies on prostate and breast cancers, metastasis-to-metastasis spread was frequent (46, 48). In our study, of 7 patients with 2 or more metastatic lesions, evidence of metastasis-to-metastasis spread was found in 2 patients (28.6%). In GB-A1 (**Figure 2A**), it appears that CBD, omentum 1-2, mesentery, and abdominal wall 2-4 lesions may originate from abdominal wall 1 (old) rather than from primary GBAC considering clone F. Similarly, in GB-A2 (**Figure 3A**), considering clinical information that mesentery and chest wall metastases occurred later than liver and lung metastases, mesentery and chest wall metastases may originate from liver and lung metastases, respectively. Although intratumoral heterogeneity of primary GBAC may make it difficult to draw a strong conclusion, our data may provide evidence of metastasis-to-metastasis spread.

Of the 11 expanded subclones at the metastasis stage, we described putative driver mutations in tumor suppressor genes in 8 subclones based on the previous literature (Table 2). For example, *SMAD4* mutations expanded during metastasis in GB-A1 and GB-S3 (**Figure 2A, B, and E**) and have been associated with distant metastasis and poor prognosis in various cancers, including GBAC (10, 17, 27, 28). In GB-S1 (**Figure 2C**), clone B containing *SLIT3* F843I evolves into E during metastasis by acquiring *ROBO1* P1360Q. The SLIT/ROBO pathway suppresses tumor progression by regulating invasion, migration, and apoptosis (29). In GB-A2 (**Figure 3A**-**C**), *DICER1* T519A is found in lung and chest wall metastases. The *DICER1* gene is associated with pleuropulmonary blastoma in children (32). In GB-S5 (**Figure 3G**), *KIAA0100* F5S and *CSMD2* E411K are found during metastasis. *KIAA0100* and *CSMD2* are frequently mutated during metastasis in adrenocortical carcinoma (34).

In mutational signature analysis, we identified six dominant mutational signatures: 1 (age), 3 (DNA double-strand break-repair), 7 (ultraviolet), 13 (APOBEC), 22 (aristolochic acid), and 24 (aflatoxin). Among them, signatures 1, 3, and 13 were commonly found in MSKCC and Shanghai datasets (**Figure 4A**) (10, 11), and have also been reported in other studies (6–9). However, these previous studies merely analyzed primary tumors and did not include metastatic tumors while our study analyzed both as well as precancerous BilIN. In our study, the limited number of SNVs and small indels per sample (median 61, range 12–241) made it difficult to compare among individual tumors (**Figure 4—figure supplement 1**). Therefore, we classified the entire mutations according to (1) the developmental stages of cancer (**Figure 4B**) and (2) the timing of development during clonal evolution (**Figure 4C**). Considering the evolutionary trajectories in cancers, we suggest that the latter criteria would be better than the former in classifying mutations according to each step of carcinogenesis and metastasis. Our data indicate that the importance of signatures 1 and 13 decreased during metastasis while the roles of signatures 22 and 24 were relatively highlighted. However, due to a lack of clinical information, we could not identify whether patients were exposed to aristolochic acid or aflatoxin.

In this cancer precision medicine era, targeted sequencing data of a single specimen are not enough to determine whether the detected mutations are clonal or subclonal. This proof- of-concept study may enable us to deeply understand the clonal evolution in GBAC. Moreover, we found that some of the mutations were clonal while a substantial proportion was subclonal, which is usually not an effective druggable target. Therefore, we believe that our study highlights the importance of precise genomic analysis of multi-regional and longitudinal samples in individual cancer patients. However, one caveat is that we cannot easily apply this to real-world patients because multi-regional and longitudinal tumor biopsies may not be feasible in most patients unless they underwent surgery and repeated biopsies. Recent studies using circulating tumor DNA have shown the possibility to easily detect mutations involved in cancer development and progression (49). By detecting clonal mutations from the carcinogenesis stage in healthy individuals, we can diagnose GBAC at an early stage. In addition, by detecting subclonal mutations in patients with GBAC, we can monitor the expanding subclones during follow-up, which enables us to detect cancer progression earlier.

This study has several limitations. First, it is not possible to obtain samples through frequent biopsy whenever desired. Thus, tumor samples were not acquired according to their developmental sequence. Second, due to intratumoral heterogeneity, the clonal composition of a small piece of the tumor may not reflect that of the entire lesion (50). Third, as the number of analyzed samples was different among patients, an accessibility bias was inevitable – the more samples the patient has, the more clones we can identify.

In conclusion, subclonal diversity developed early in precancerous lesions and the clonal selection was a common event during malignant transformation in GBAC. However, cancer clones continued to evolve in metastatic tumors and thus maintained subclonal diversity. Our novel approach may help us to understand the GBAC of individual patients and to move forward to precision medicine that enables early detection of carcinogenesis and metastasis, and effective targeted therapy in these patients.

## Methods

### Patients and tumor samples

Patients were eligible for this study if they were diagnosed with GBAC and received surgery between 2013 and 2018 at Seoul National University Bundang Hospital (SNUBH), Seongnam, Korea. Among 11 enrolled patients, two patients underwent rapid autopsy after death. Patients’ clinical information was obtained through retrospective medical record reviews. This study was approved by the institutional review board (IRB) of SNUBH (IRB No. B-1902/522-303).

### Whole exome sequencing (WES) of GBAC

DNA was extracted from fresh-frozen or FFPE tissues. Library preparation and exome capture was carried out using Agilent SureSelect^XT^ Human All Exon V6 (Santa Clara, CA, USA). WES was conducted with a paired-end, 100-bp using Illumina NovaSeq 6000 (San Diego, CA, USA). The depth of coverage of tumors and normal control samples were at least 300× and 200×, respectively.

### Analysis of WES data

WES data of GBAC and matched normal samples were analyzed using the Genomon2 pipeline (Institute of Medical Science, University of Tokyo, Tokyo, Japan; https://genomon.readthedocs.io/ja/latest/, accessed on Feb. 1, 2022) as previously described (39). In brief, sequencing reads from adapter-trimmed .fastq files were aligned to the human reference genome GRCh37 (hg19) without the ‘chr’ prefix using Burrows-Wheeler Aligner version 0.7.12, with default settings. Somatic single nucleotide variants (SNVs) and small indels were called by eliminating polymorphisms and sequencing errors and filtered by pre-specified criteria: (a) only exonic or splicing sites were included; (b) synonymous SNVs, unknown variants, or those without proper annotation were excluded; (c) polymorphisms in dbSNP 131 were excluded; (d) p-values < 0.01 from Fisher’s exact test were included; (e) simple repeat sequences were excluded; (f) strand ratio between positive-strand and negative-strand should not be 0 or 1 in tumor samples; (g) the number of variant reads should exceed 4 in tumor samples. For each patient, filtered variant lists of tumor samples were merged. Then, the merged list of target variants was manually called in each .bam file using bam-readcount version 0.8.0 (https://github.com/genome/bam-readcount, accessed on Feb. 1, 2022) with Phred score and mapping quality of more than 30 and 60, respectively.

For samples with the tumor purity > 0.4, the ploidy of tumor cells was estimated using Sequenza version 3.0.0 to identify whole genome doubling (WGD) (21). Copy number variations (CNVs) were analyzed using the Control-FREEC version 11.5 (51). In brief, the aligned .bam files were converted to .pileup.gz format using SAMtools version 1.9 (52). The .pileup.gz files of data from GBAC and matched normal samples were analyzed using Control-FREEC with default settings. These datasets were then used to statistically infer clonal population structure using PyClone version 0.13.0 with the ‘pyclone_beta_binomial’ model (22). Clonal phylogeny was inferred by CITUP version 0.1.0 (23) using cellular prevalence values of each cluster which were generated by PyClone. To ensure accurate tree construction, clusters containing only one mutation were excluded from the input to CITUP if the mutation’s role is unclear from previous literature. This filter removed 42 mutations from a total of 1,577 mutations, representing less than 2.7% of all clustered mutations. In addition, in GB-A2, clusters which limited to only one organ were excluded from the analysis because the calculation for the phylogenetic tree using CITUP took more than one month when the number of clusters was ≥ 14. Analyzed results were visualized using MapScape for multi-regional specimens and TimeScape for longitudinal or putative longitudinal datasets, as appropriate (24). In all analysis steps, the data were adjusted for the tumor purity values of each tumor.

The mutational signatures of SNVs were analyzed using Mutalisk (53). To compare with the other GBAC cohorts, we additionally analyzed two public datasets of GBAC from the Memorial Sloan Kettering Cancer Center (MSKCC) and the Shanghai group (10, 11), which could be downloaded from the cBioPortal (https://www.cbioportal.org/, accessed on Feb. 1, 2022) (54).

### *HER2* immunohistochemistry (IHC) and silver in situ hybridization (SISH)

The histologic sections from individual FFPE tissues were deparaffinized and dehydrated. IHC and SISH analysis of *HER2*-positive cells was conducted by the board-certified pathologist using PATHWAY anti-*HER2*/neu antibody (4B5; rabbit monoclonal; Ventana Medical Systems, Tucson, AZ, USA) and a staining device (BenchMark XT, Ventana Medical Systems, Tuscon, AZ, USA), respectively, as previously described (55). Signals from 20 tumor cells were counted and a *HER2*/CEP17 ratio ≥ 2.0 was defined as *HER2* amplification (19). Wilcoxon rank-sum test is used to compare the mean *HER2*/CEP17 ratio of BilIN versus primary tumor.

## Acknowledgments

This study was supported by a grant from Seoul National University Bundang Hospital Research Fund (No. 16-2021-001) and the Small Grant for Exploratory Research (SGER) program (NRF-2018R1D1A1A02086240) of National Research Foundation (NRF), Korea. The authors would like to express the deepest respect to the two patients who donated their bodies for this study after death.

## Competing interests statement

All other authors declare no competing interests.

## Data availability statement

The raw sequence data underlying this manuscript are available as fastq files at the NCBI SRA database under the BioProject number PRJNA821382.

---

**Figure 1—figure supplement 1.**
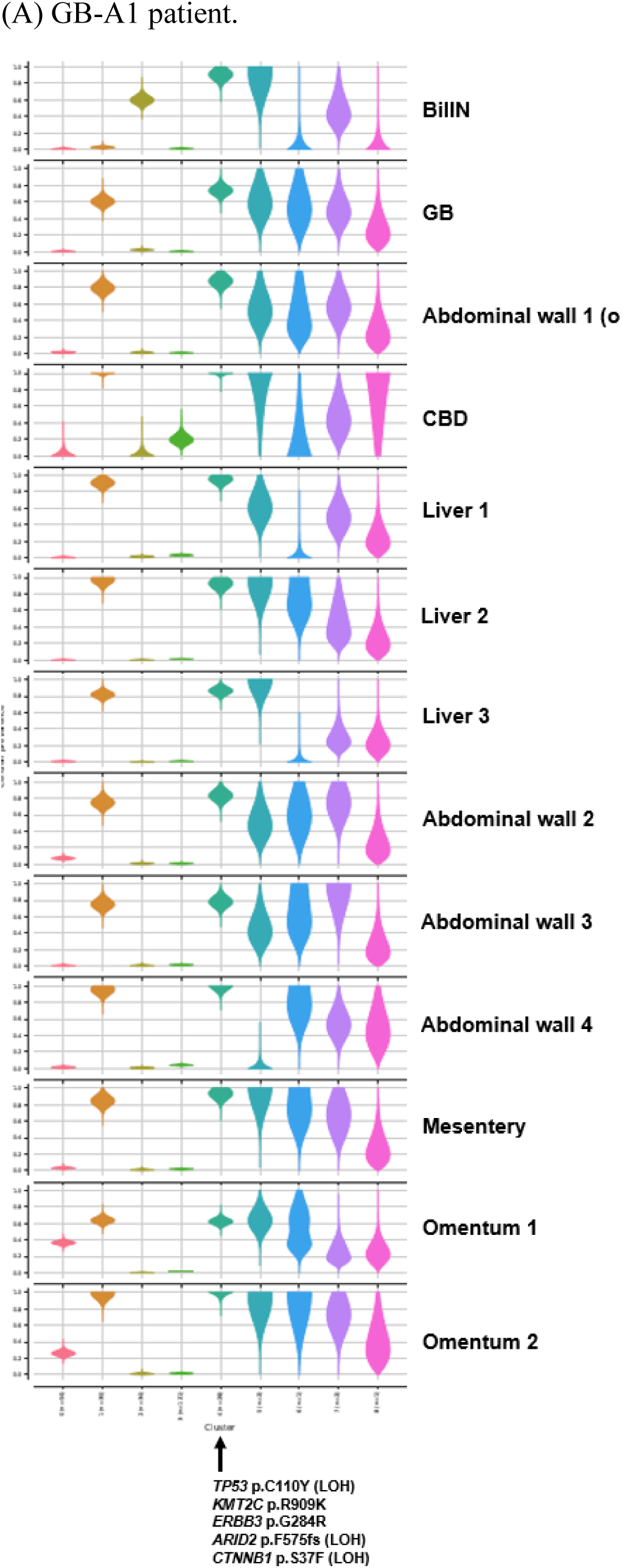
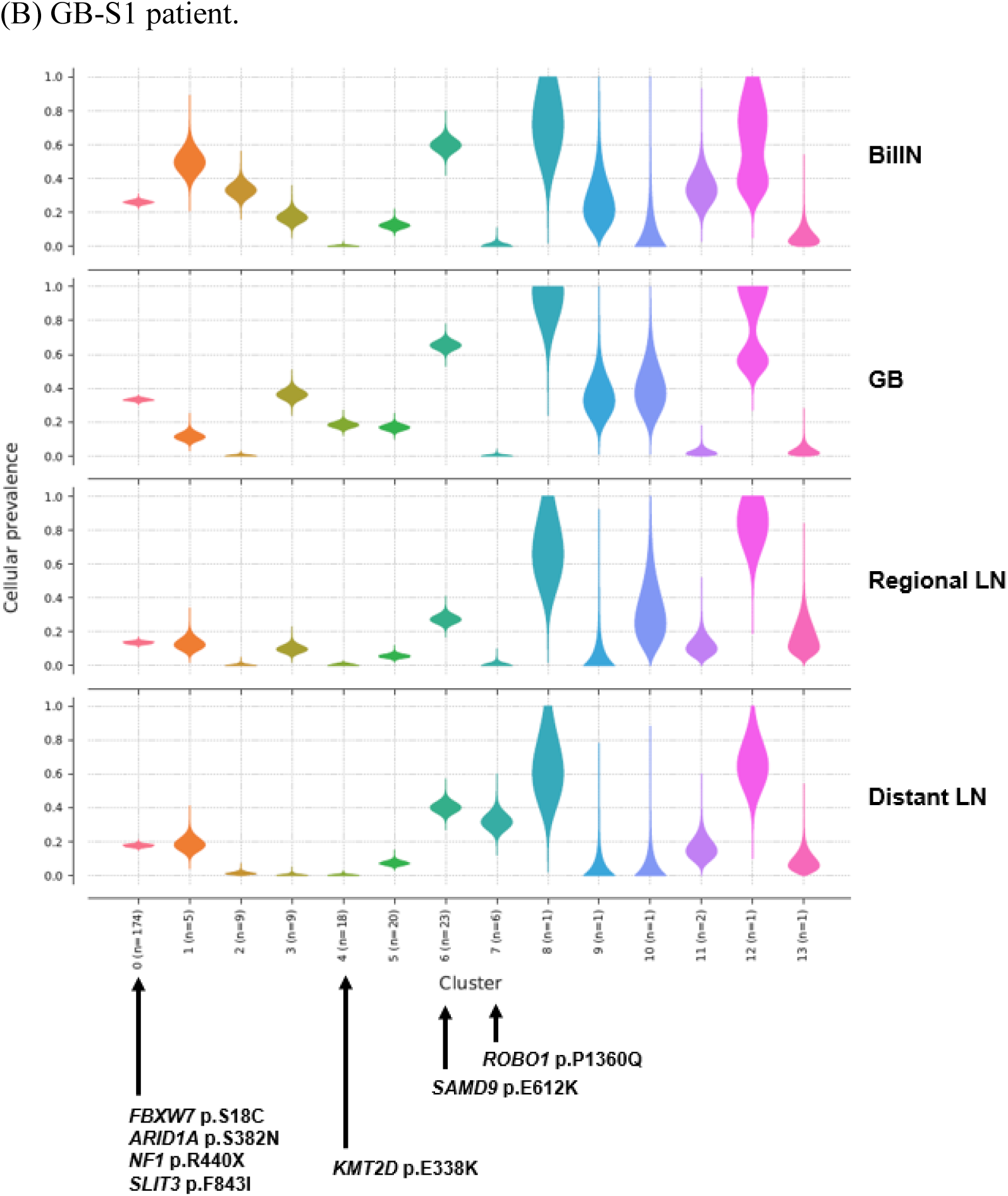
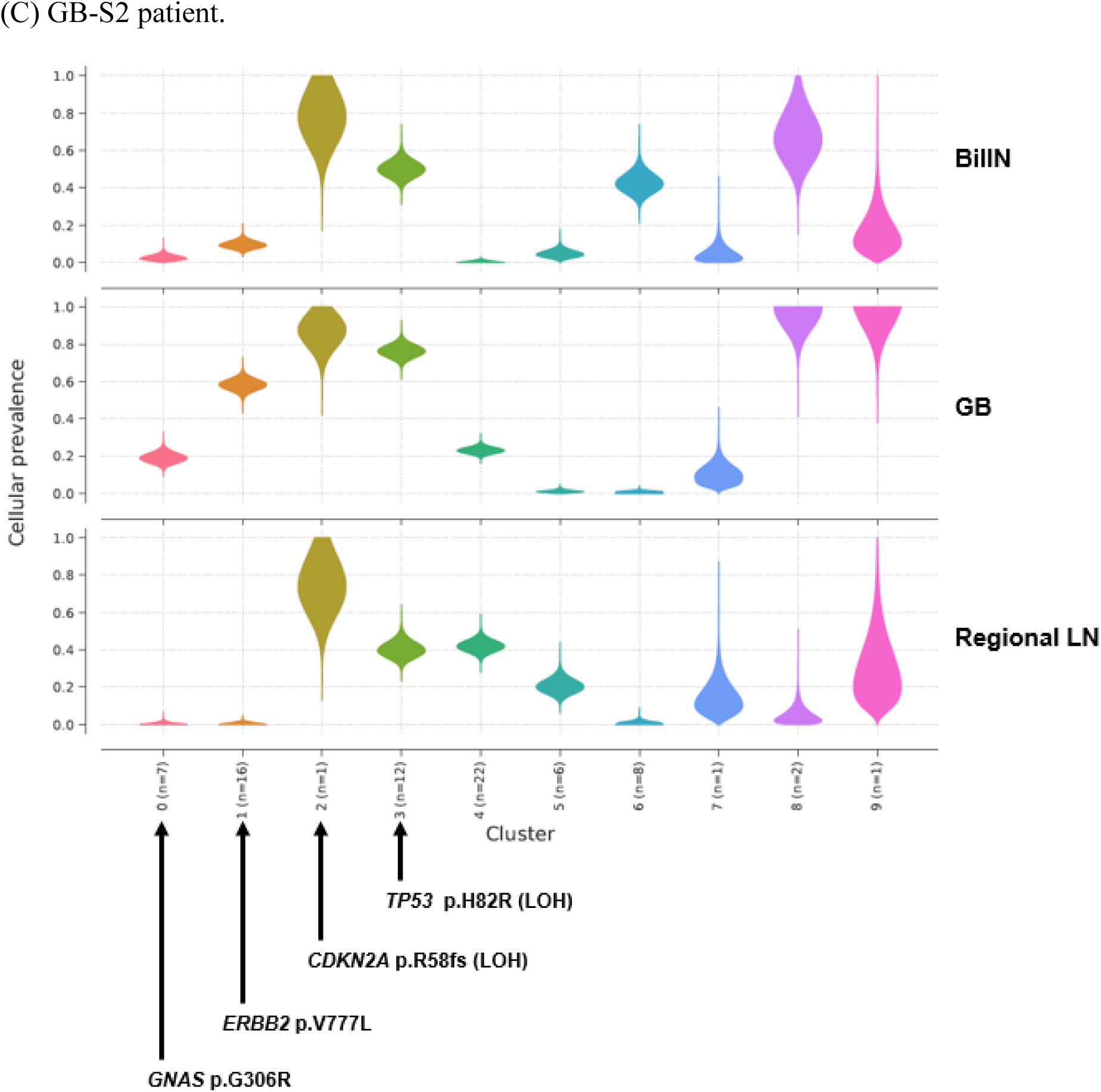
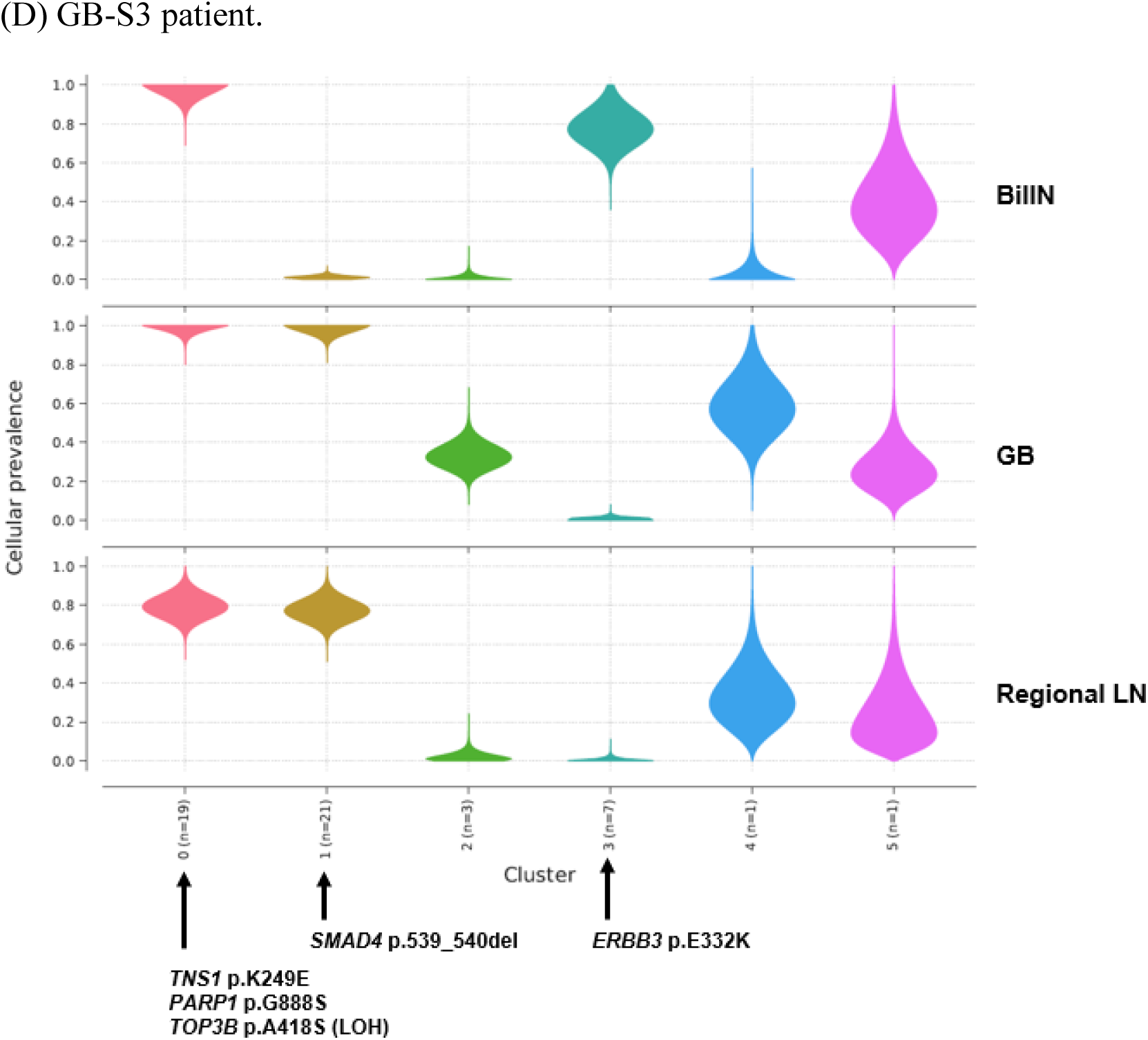
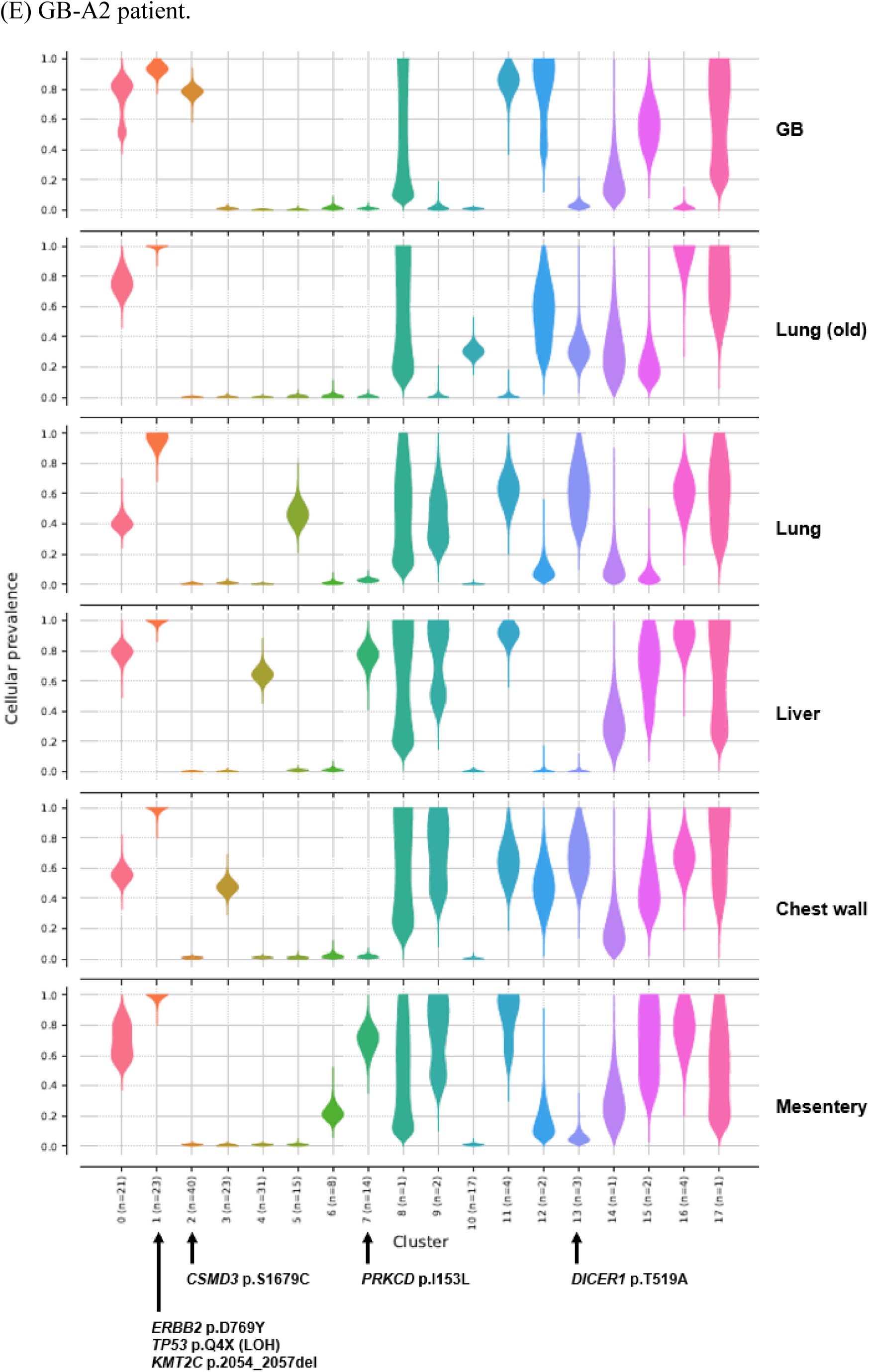
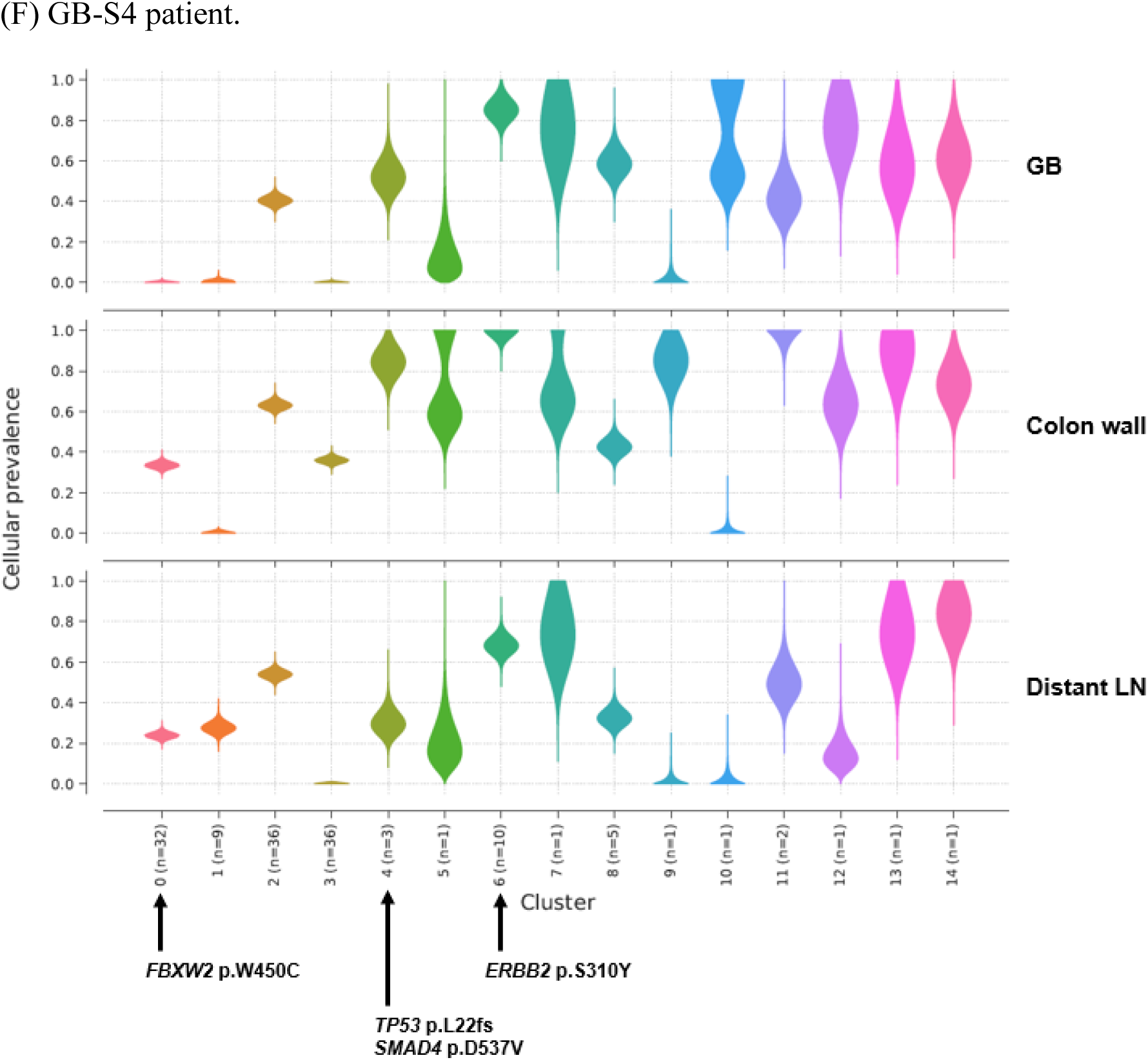
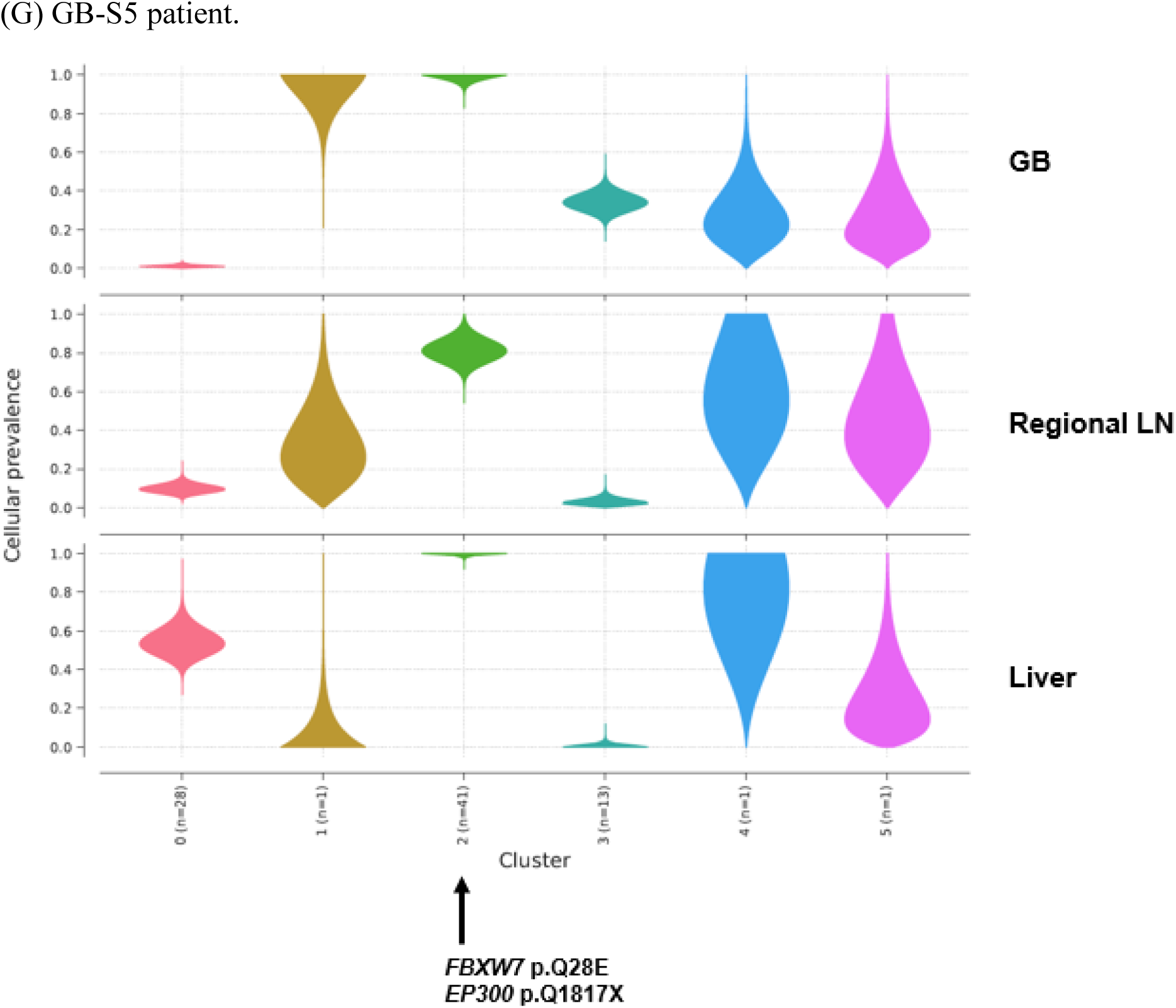
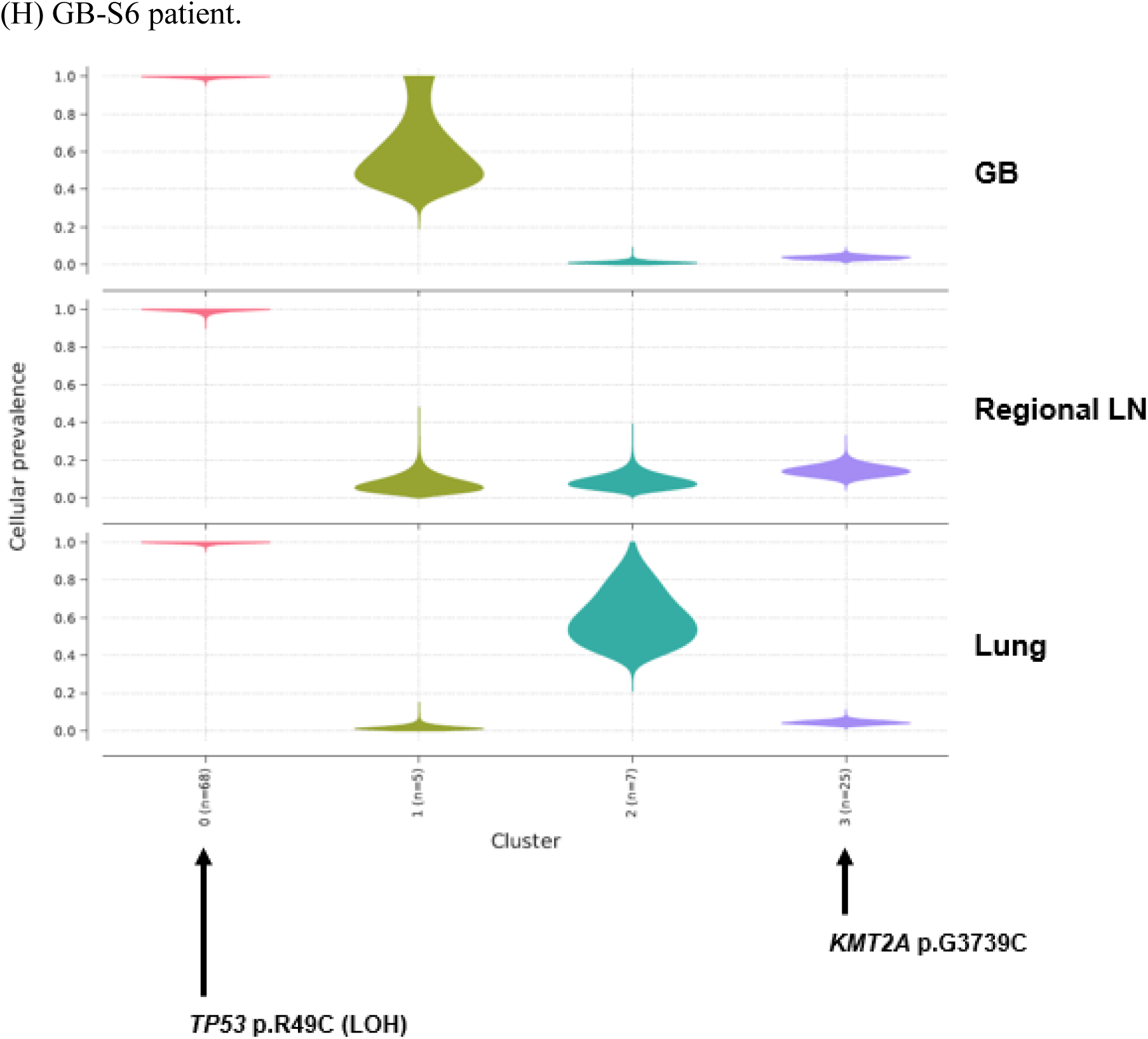
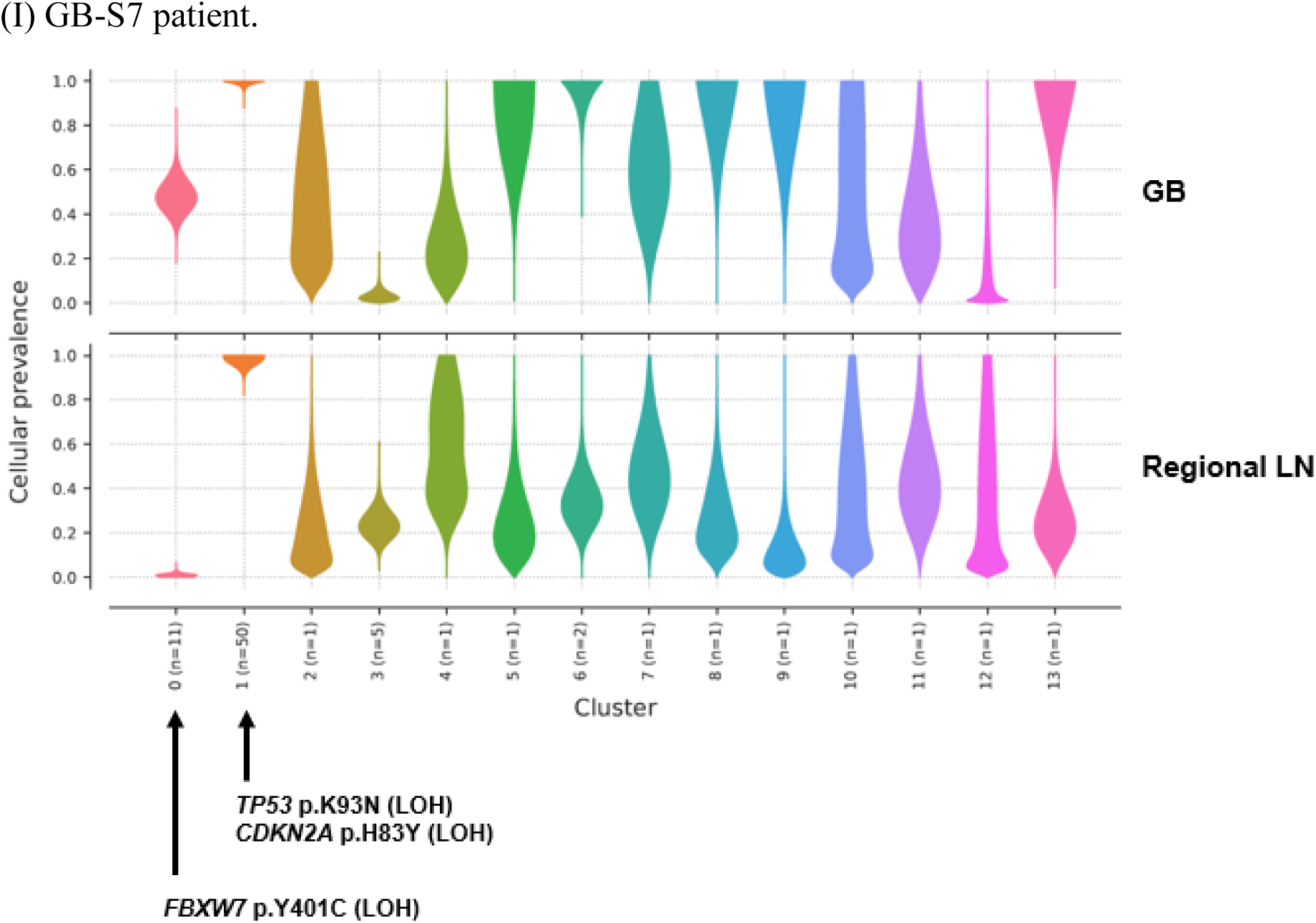
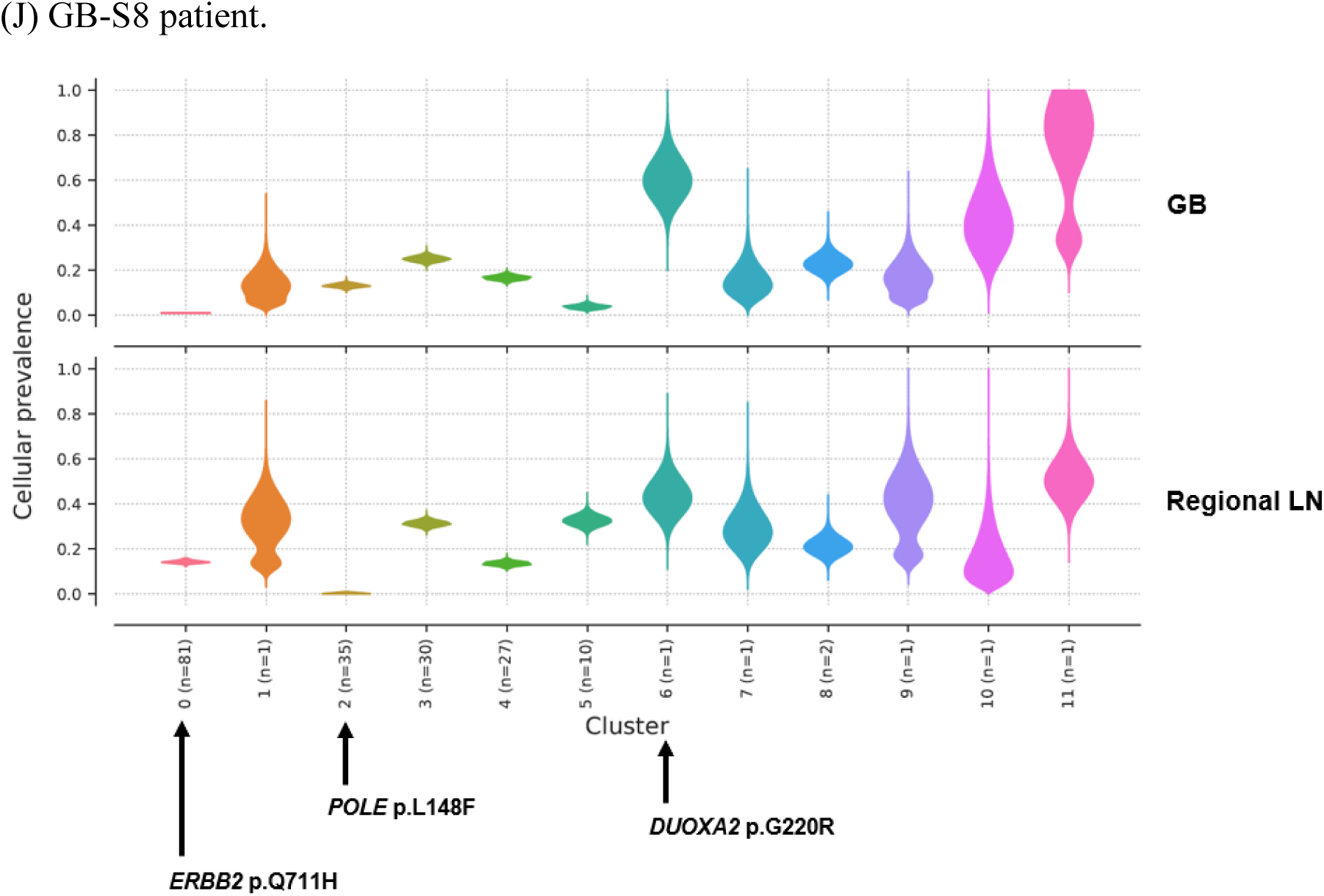
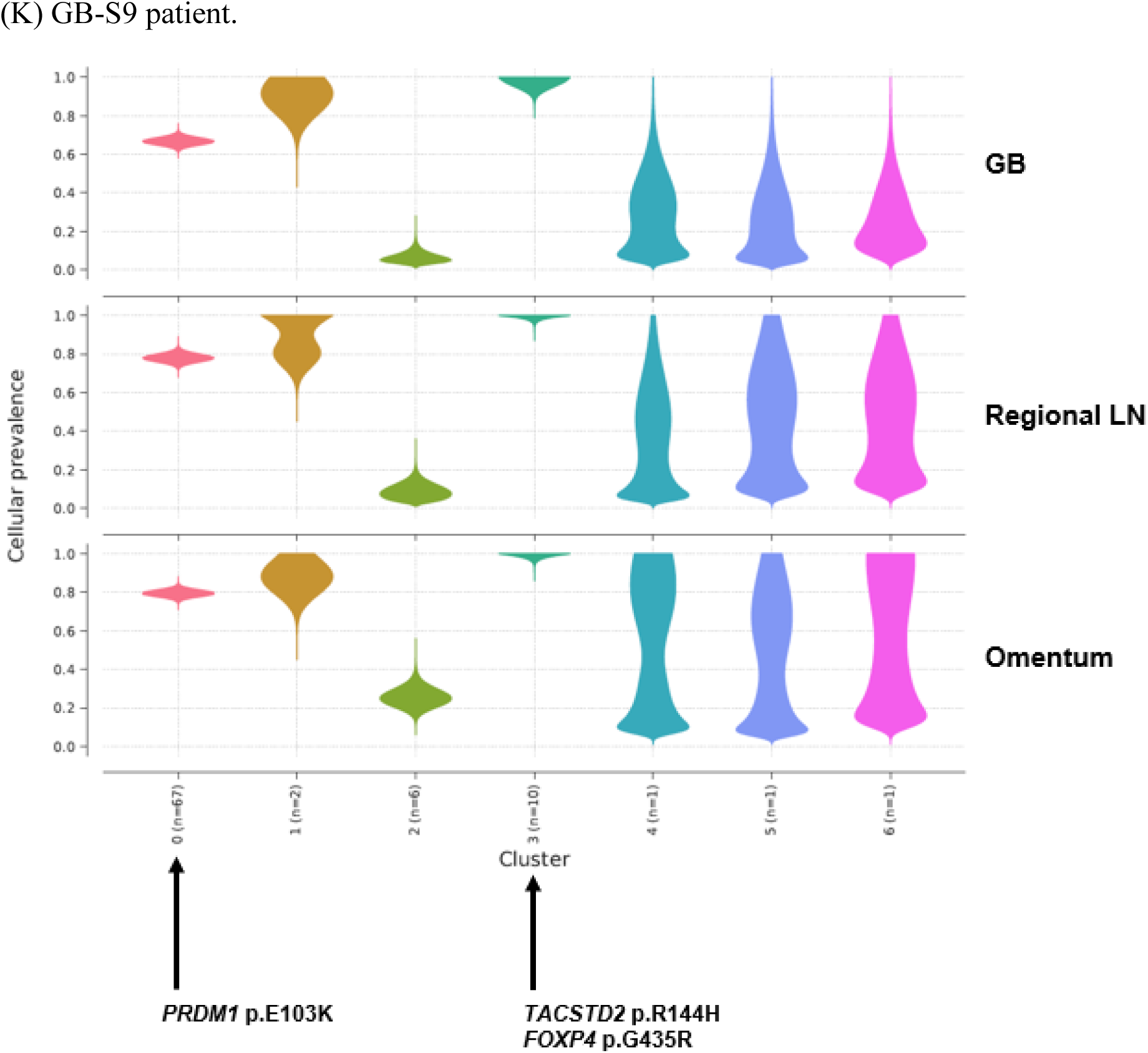
Inference of clonal structure by PyClone algorithm. **(A-K)** The Clonal population structure was statistically inferred for GB-A1 **(A),** GB-S1 **(B)**, GB-S2 **(C)**, GB-S3 **(D)**, GB-A2 **(E)**, GB-S4 **(F)**, GB-S5 **(G)**, GB-S6 **(H)**, GB-S7 **(I)**, GB-S8 **(J)**, and GB-S9 **(K)** using PyClone. BilIN, biliary intraepithelial neoplasia; GB, gallbladder; CBD, common bile duct; LN, lymph node.

**Figure 1—figure supplement 2.**
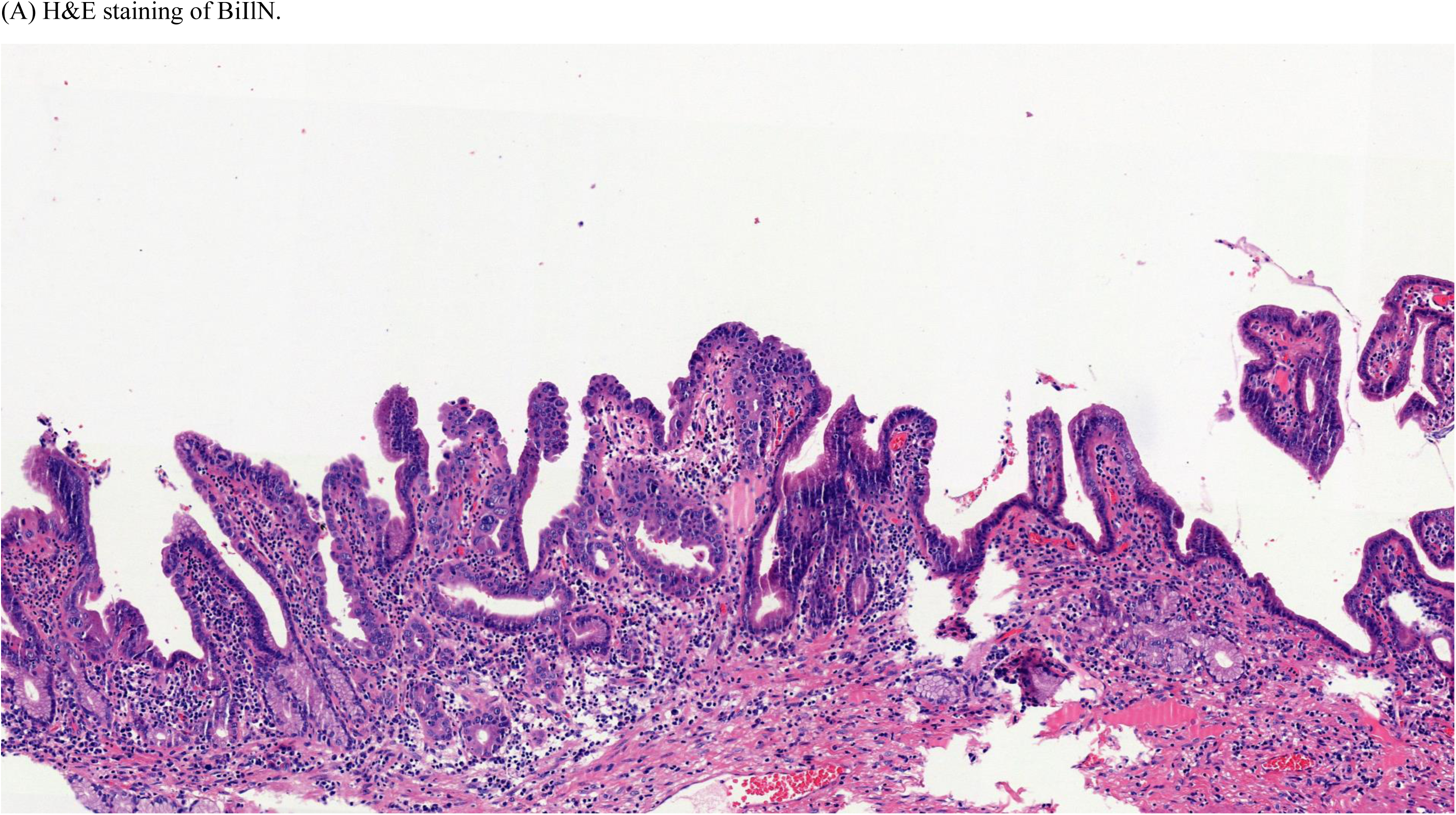
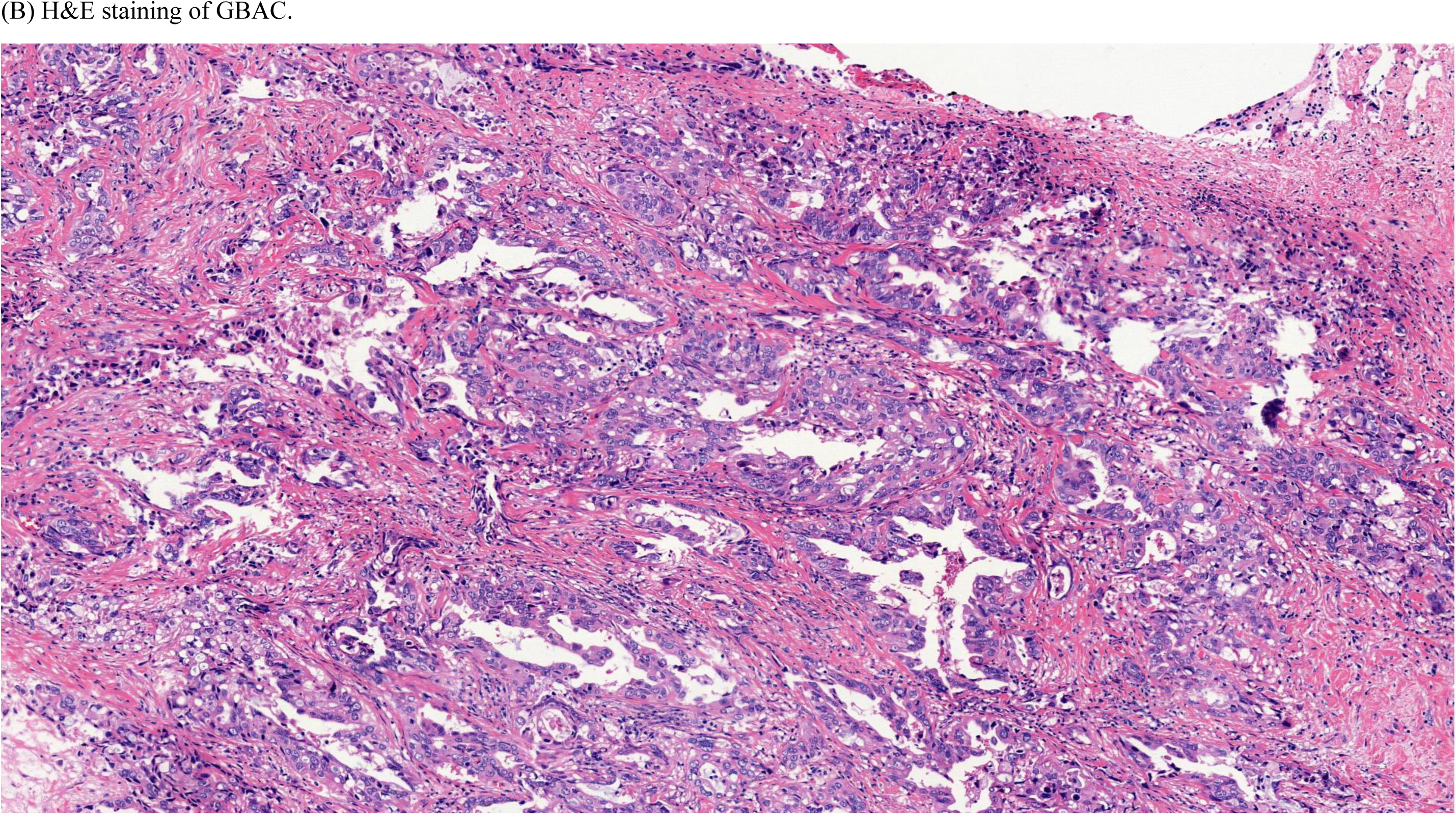
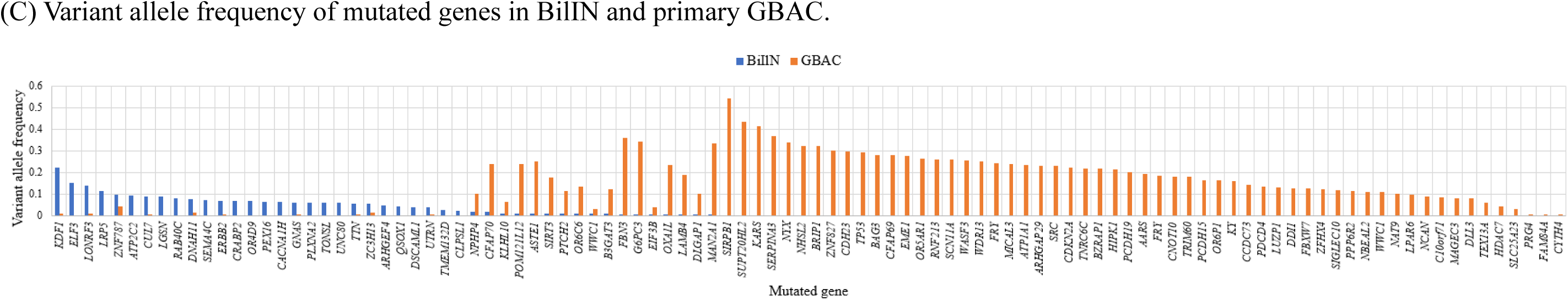
BiIlN and primary GBAC of the GB-S7 patient presumed to be derived from different origins. **(A)** H&E staining of BiIlN. **(B)** H&E staining of GBAC. **(C)** Variant allele frequency of mutated genes in BilIN and primary GBAC. BilIN, biliary intraepithelial neoplasia; H&E, hematoxylin, and eosin.

**Figure 4—figure supplement 1.**
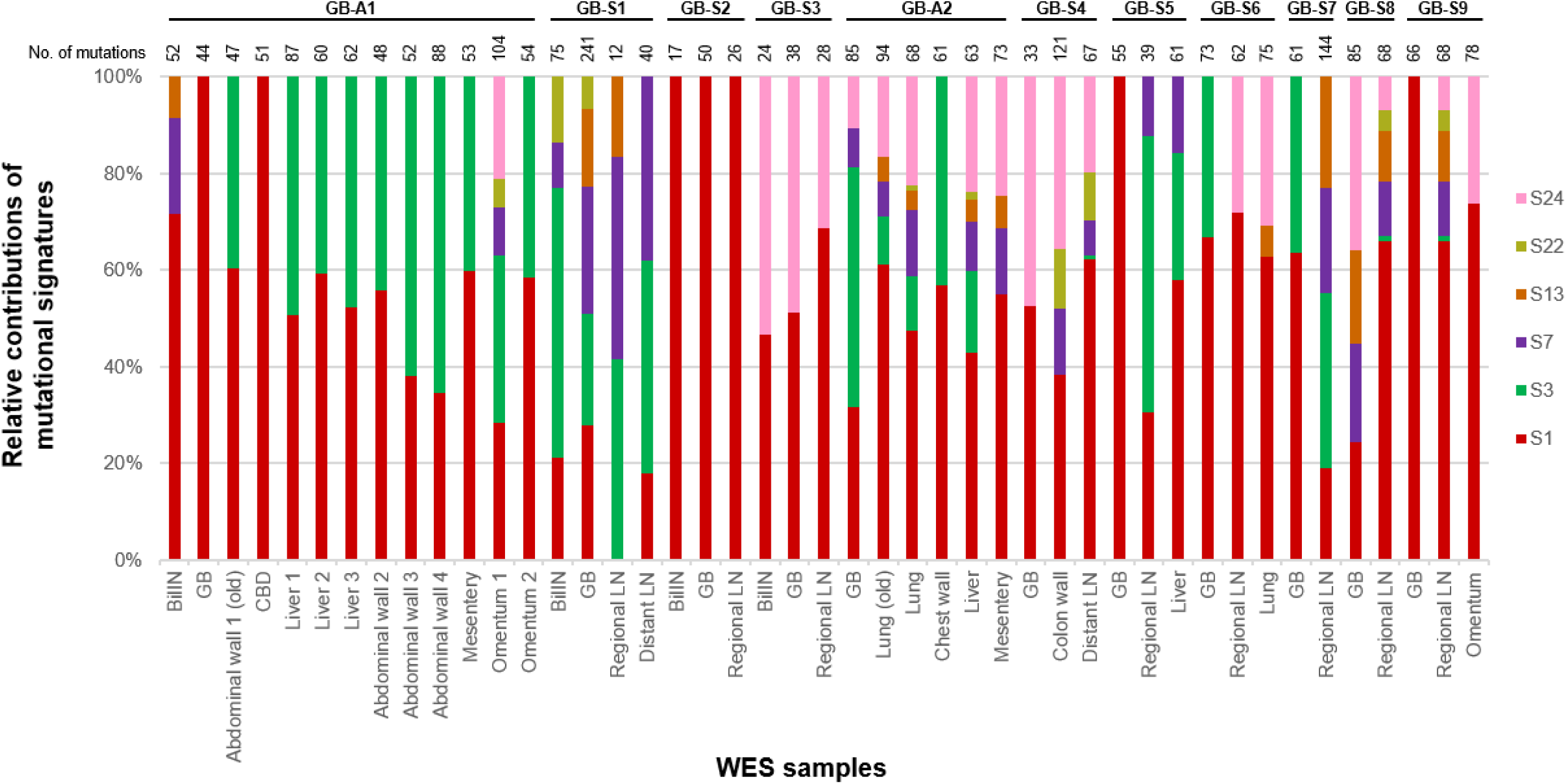
The 100% stacked bar plots comparing the proportions of known COSMIC mutational signatures within each sample from 11 GBAC patients. BilIN, biliary intraepithelial neoplasia; COSMIC, catalogue of somatic mutations in cancer; GB, gallbladder; LN, lymph node.

## Notes

### Competing Interest Statement

The authors have declared no competing interest.

